# Heme orchestrates a tissue stress response to proteolytic damage

**DOI:** 10.64898/2026.05.31.729126

**Authors:** Karen Agaronyan, Allison M. Greaney, Shuang Yu, Arun R. Chavan, Darine W. El-Naccache, Edward P. Manning, Lokesh Sharma, Charles S. Dela Cruz, Ruslan Medzhitov

**Affiliations:** Department of Immunobiology, Yale University; New Haven, CT, USA; Section of Pulmonary, Critical Care, and Sleep Medicine, Yale University; New Haven, CT, USA; VA Connecticut Healthcare System; West Haven, CT, USA; Division of Pulmonary, Allergy, Critical Care and Sleep Medicine, University of Pittsburgh; Pittsburgh, PA, USA; Section of Pulmonary, Critical Care and Sleep Medicine, Yale University; New Haven, CT, USA; Howard Hughes Medical Institute, Department of Immunobiology, and Tananbaum Center for Theoretical Biology, Yale University; New Haven, CT, USA

## Abstract

Whereas cellular stress responses are well defined, tissue-level stress remains poorly understood. Proteases are among the most widespread enzymes, and excessive proteolytic activity drives diseases such as arthritis and chronic obstructive pulmonary disease, yet unifying features of this stress are unclear. Here, using the lung and diverse proteases, we identify a conserved injury signature of proteolytic stress marked by vascular disruption, red blood cell extravasation, and heme release that triggers oxidative stress. We show that alveolar macrophages act as primary sensors of this stress response, activating NRF2-dependent heme detoxification program and fibroblasts produce protease inhibitors to limit damage. Repeated exposure to proteolytic stress induces tissue adaptation and protects against subsequent injury and infection. These findings define a unifying framework for tissue-level proteolytic stress sensing and adaptation.

---

Animals continuously monitor and regulate homeostatic variables such as blood glucose concentration, temperature, pH and oxygen tension in order to preserve function and survival. When these variables deviate significantly from their set points, stress-response programs are engaged to restore homeostasis (*1–3*). At the cellular level, these responses are well characterized including hypoxia and endoplasmic reticulum (ER) stress, which are sensed by dedicated pathways and transcriptional regulators that act to mitigate oxygen deprivation or proteostatic imbalance, respectively (*4, 5*). By contrast, stress responses at the tissue level remain poorly defined, despite their importance in maintenance of tissue homeostasis. Moreover, tissue-level perturbations cause structural and functional damage that underlies inflammatory diseases such as inflammatory bowel disease, acute respiratory distress syndrome, and chronic wounds (*6–8*). Understanding how tissues sense and respond to stress and damage is critical both for biology of homeostasis and for designing interventions that promote tissue resilience and repair.

Proteolytic damage is a widespread, non-infectious source of macromolecular injury in tissues. Proteases derived from endogenous sources, microbial toxins, venoms, and industrial formulations are among the most abundant enzymes and potent mediators of tissue damage (*9–11*). Excessive proteolytic activity contributes to and drives disease states such as arthritis, chronic obstructive pulmonary disease (COPD), nonhealing wounds, and cancer (*12–15*). Despite its prevalence, the unifying features of proteolytic tissue damage are not well establishe d with reported consequences ranging from hemorrhagic necrosis, basement-membrane disruption, and cartilage degradation (*16–18*). Resident immune cells, including macrophages and mast cells, as well as nonimmune cells such as epithelial stem cells, can serve as sensors of tissue stress (*3, 19–22*). However, the specific cellular sensors that detect proteolytic tissue stress and the circuits they engage in remain poorly defined. Here, using a single tissue (lung) and a panel of proteases with distinct catalytic activities and origins, we show that proteolytic stress produces a conserved injury signature characterized by vascular disruption, red blood cell extravasation and heme release, which drives a tissue oxidative stress response. We identify alveolar macrophages (AMs) as the principal sensors of heme-driven oxidative stress that activate the heme detoxification program, and fibroblasts as primary producers of protease inhibitors that limit ongoing proteolytic activity. Together, these findings define a unifying feature of proteolytic damage and delineate the molecular and cellular circuits that respond to proteolytic stress in lung tissue. More broadly, they provide a new framework for understanding how tissues detect and mitigate macromolecular injury and highlight potential targets for limiting tissue damage.

## Results

### Unifying features of proteolytic tissue stress

To define unifying features of proteolytic tissue stress, we employed three exogenous proteases representing the major catalytic classes that account for ∼90% of known protease activities (*9, 10*): the cysteine protease papain (*Carica papaya*), the serine protease subtilisin (*Bacillus subtilis*), and the Zn-dependent metalloprotease LasB (*Pseudomonas aeruginosa*). Our previous studies have validated that these proteases are enzymatically active, endotoxin-free, and do not activate pattern recognition receptors. Previously, we also established that *in vivo* effects of these proteases depend on their catalytic activities (*23–26*). Because exogenous proteases at high concentrations exhibit promiscuous substrate specificities, they provide an ideal model for inducing generalized proteolytic stress (*27, 28*).

Lung is continuously exposed to environmental proteolytic stress during infections and sterile inflammation in which protease-antiprotease imbalance contributes to diseases such as common obstructive pulmonary disease (COPD) and a1 -antitrypsin deficiency (*13, 29–31*). To develop a model of proteolytic tissue stress, we administered proteases intranasally either once (labelled 1d) or three times at 24-hour intervals (labelled 3d), thus modeling acute and repeated stress, respectively (Fig. 1A, top). The dose per administration remained consistent across groups, meaning that 3d animals received threefold higher cumulative exposure. We measured experimental parameters 24 hours after the final challenge for papain and subtilisin and 48 hours for LasB (Fig. 1A, top). A single exposure (1d) to each protease resulted in increased protein levels in bronchoalveolar lavage (BAL), red blood cell (RBC) infiltration, and platelet accumulation (Fig. 1A-C). Consistent with vascular leakage and hemorrhage, BAL from 1d animals contained elevated hemoglobin, hematocrit, and free heme (Fig. 1D-F). These findings identify vascular injury with hemoglobin and heme release as a unifying hallmark of proteolytic tissue stress across distinct classes of proteases. Remarkably, animals exposed repeatedly (3d) exhibited markedly attenuated responses, comparable to PBS-treated controls (Fig. 1A-F), indicating the development of tissue adaptation that protects from repeated challenges.

**Fig. 1.**
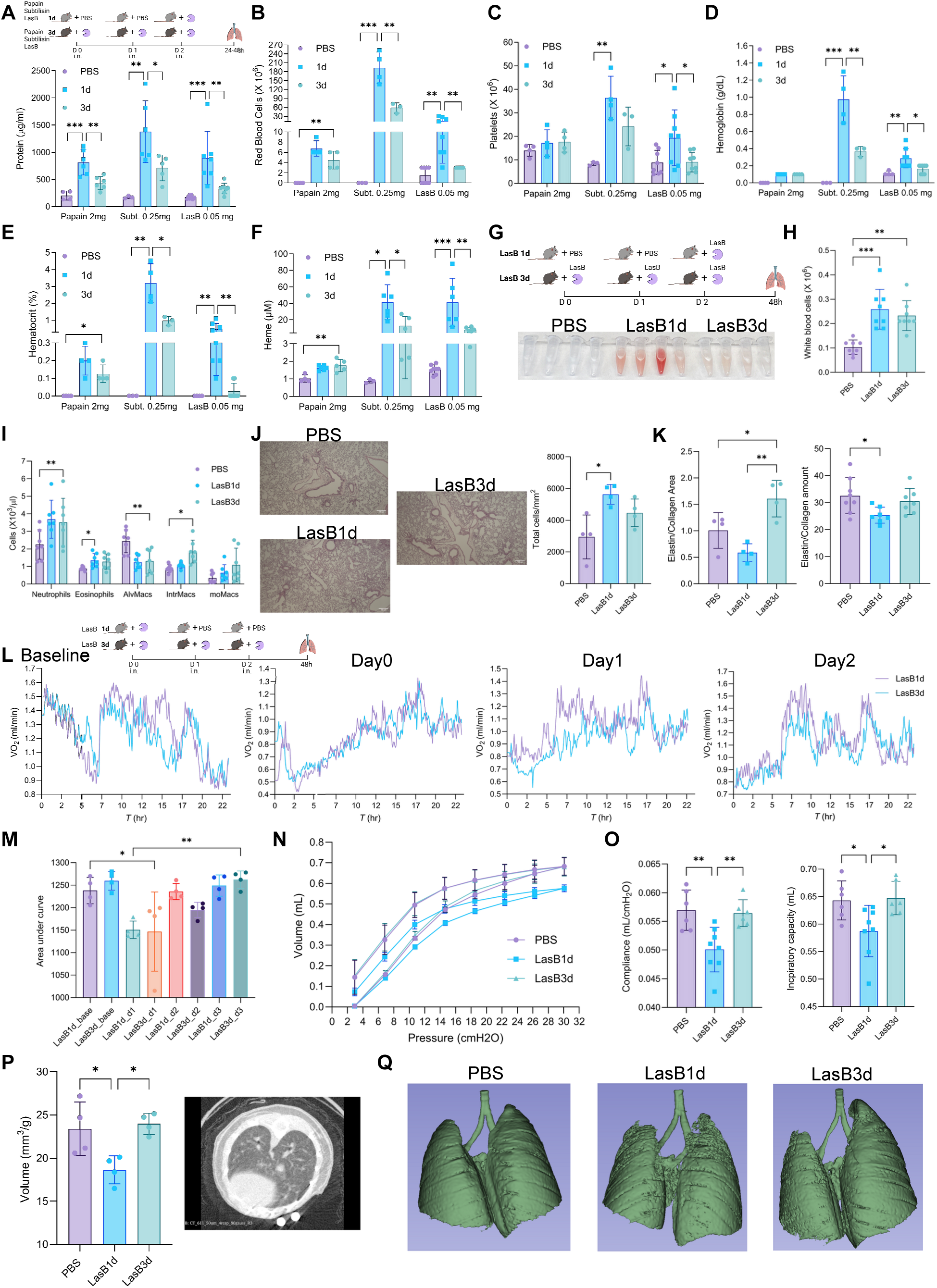
Adaptation to proteolytic tissue stress. **(A-F)** Experimental model of proteolytic tissue stress (A, top), BAL protein levels (A, bottom), RBC (B) and platelet (C) counts, hemoglobin (D), hematocrit (E) and heme (F) levels after 1d or 3d exposure to proteases (n = 3-8 mice). **(G)** Experimental model of LasB tissue stress (top), photograph of BAL after LasB treatment (bottom). **(H-I)** BAL WBC (H) and lung myeloid cell (I) numbers after acute and repeated LasB exposure (AlvMacs – alveolar macrophages, IntrMacs – interstitial macrophages, moMacs – monocyte-derived macrophages (n = 7-8 mice). **(J)** Representative lung images of H&E staining (left) and quantification (right) after acute and repeated LasB exposure (n = 4 mice). **(K)** Ratio of elastin to collagen based on trichrome and EVG staining (left) and biochemical analysis (right) acute and repeated LasB exposure (n = 4-8 mice). **(L-M)** Experimental model of modified LasB protocol (L, top left), VO^2^ curves (L) and area under the curve of respiratory exchange ratio (RER) (M) during acute and repeated LasB exposures (n = 4 mice). **(N-O)** Pressure-volume loops (N), respiratory compliance (O, left) and inspiratory capacity (O, right) acute and repeated LasB exposure (n = 6-8 mice). **(P-Q)** µCT lung volume measurements (P left), representative µCT image of lung (P right) and µCT-based 3D reconstructions of lungs acute and repeated LasB exposure (n = 4 mice). Data are mean +/- SD and were analyzed by one-way ANOVA with Bonferroni multiple comparison test. (*p=0.05, **p=0.01, ***p=0.001).

Among the proteases tested, the bacterial metalloprotease LasB displayed the greatest potency. A dose of 0.05 mg/kg LasB induced levels of BAL protein, RBC and platelet infiltration, vascular damage, and heme release comparable to 0.25 mg/kg subtilisin or 2 mg/kg papain (Fig. 1A-F), corresponding to ∼5-fold and ∼40-fold greater potency, respectively. Increasing the LasB dose to 0.08 mg/kg further amplified BAL protein, cellular infiltration, hemoglobin, hematocrit, and heme levels (Fig. S1A-C). Based on these data, we selected 0.05 mg/kg LasB for subsequent studies of acute (LasB1d) and repeated (LasB3d) proteolytic stress (Fig. 1G, top). Acute LasB1d challenge was visibly associated with hemorrhagic BAL (Fig. 1G bottom).

Proteolytic stress triggered robust leukocyte infiltration (Fig. 1H, S2A), characterized by increased neutrophils, interstitial macrophages, and monocyte-derived macrophages, accompanied by depletion of alveolar macrophages (AMs) (Fig. 1I, S1D). These changes parallel responses described in bacterial-, viral-, and bleomycin-induced lung injury (*32–37*). Classical monocytes, dendritic cells, T cell subsets, and innate lymphoid cells were largely unchanged (Fig. S3A-C). Consistent with vascular injury, total endothelial cells were reduced, whereas epithelial cells and fibroblasts were preserved (Fig. S3D). Histological analysis revealed increased cellular infiltration and reciprocal alterations in collagen and elastin area and total content (Fig. 1J, S2H-I). The elastin-to-collagen ratio decreased during acute 1d stress and normalized after repeated 3d exposure (Fig. 1K). BAL from LasB1d animals contained elevated IL-6, CXCL10, and CCL3 cytokines (Fig. S2B), consistent with acute inflammatory activation and recruitment of neutrophils and monocytes. Other BAL or serum cytokines were not significantly altered (Fig. S2C-D). Systemic metabolites and enzymes were unchanged except for reduced chloride levels during acute stress (Fig. S2E). LasB1d animals exhibited transient hypothermia (Fig. S4A), with trends toward reduced oxygen saturation, heart rate, and respiratory rate (Fig. S4B), all of which normalized in LasB3d animals. To characterize systemic metabolic responses, we performed metabolic cage analysis using a modified protocol in which LasB1d animals received protease on day 0 and PBS, thereafter, allowing direct temporal comparison with repeatedly challenged LasB3d animals. On day 0, both groups showed reduced VO2, VCO2, respiratory exchange ratio (RER), energy expenditure, and food and water intake (Fig. 1L, S5). On day 1, LasB1d animals receiving PBS showed no decline, whereas LasB3d animals exhibited suppression following the second protease exposure. On day 2, however, LasB3d animals challenged a third time displayed metabolic curves indistinguishable from LasB1d group receiving PBS, indicating metabolic adaptation (Fig. 1L, S5). Area under the curve analysis of RER further substantiated this adaptive response (Fig. 1M).

To investigate changes in pulmonary mechanics and lung volume, we performed pulmonary function test and µCT. LasB1d animals demonstrated markedly altered pressure-volume loops relative to LasB3d and PBS groups (Fig. 1N). Lung compliance, inspiratory capacity, and respiratory system compliance (Crs) were significantly reduced in LasB1d animals and restored in LasB3d animals (Fig. 1O, S4D). Tissue damping (G) and elastance (H) were elevated during acute stress and normalized after repeated exposure, with an increased G/H ratio in both groups (Fig. S4C-D). These findings are consistent with pulmonary edema, vascular injury, and hemorrhage during acute proteolytic stress, followed by functional recovery upon repeated exposure. µCT imaging confirmed preserved lung volumes in LasB3d animals comparable to PBS controls (Fig. 1P) and 3D lung reconstructions revealed substantial structural disruption in LasB1d lungs that was attenuated in LasB3d animals. Collectively, these results demonstrate that diverse proteolytic activities converge on a common acute response characterized by vascular injury, hemorrhage, and heme release, accompanied by impaired metabolic and pulmonary function. Strikingly, repeated proteolytic exposure induces broad tissue adaptation, restoring vascular damage, metabolic homeostasis, and lung mechanics despite continued enzymatic challenge.

### Anti-oxidative transcription factor NRF2 is necessary for adaptation to proteolytic tissue stress

To elucidate the mechanisms underlying tissue adaptation to proteolytic stress, we combined genetic and pharmacological approaches to interrogate candidate cellular and molecular pathways. Several immune populations including eosinophils, neutrophils, and monocytes, have been implicated in driving tissue injury during infectious and sterile inflammation (*35, 36, 38–40*). Consistent with this, we observed increased frequencies of eosinophils, neutrophils, and monocytes during proteolytic stress (Fig. 1I). To determine the contribution of eosinophils, we used two complementary depletion strategies. Administration of an a-CCR3 antibody effectively reduced eosinophil numbers (Fig. S6A) but did not alter BAL protein levels during repeated proteolytic challenge (Fig. 2A). Similarly, ΔdblGata-deficeint (KO) mice, which lack eosinophils (*41*) (Fig. S6C), exhibited comparable BAL protein and RBC levels during tissue adaptation (Fig. S6B). These findings indicate that eosinophils are dispensable for adaptation to proteolytic stress. Neutrophil depletion using a-Ly6G antibody resulted in their reduction (Fig. S6D) without affecting tissue adaptation. Platelets were also increased in BAL during proteolytic stress (Fig. 1C) and given their established roles in hemostasis and tissue repair (*42, 43*), we tested their functional contribution using clopidogrel to inhibit platelet aggregation. Clopidogrel treatment did not alter BAL protein, platelet or RBC counts, nor granulocyte and macrophage composition (Fig. S6F-G), indicating that platelet aggregation is not necessary for adaptation. Recent studies have highlighted critical roles for recruited monocytes and monocyte -derived macrophages in tissue repair following depletion of resident macrophages in infectious and sterile lung injury(*35, 36, 38*). Consistent with this, proteolytic stress resulted in depletion of AMs and expansion of interstitial and monocyte-derived macrophages (Fig. 1I). To assess the contribution of recruited monocytes, we compared WT and CCR2 KO mice, as CCR2 is the principal chemokine receptor mediating monocyte recruitment (*44*). CCR2 KO exhibited reduced interstitial and monocyte-derived macrophage populations (Fig. S7B), yet showed levels of BAL protein, RBCs, platelets, hemoglobin, hematocrit, and heme comparable to WT controls during repeated stress (Fig. 2B, S7A, C-D). Thus, recruited monocytes are not necessary for tissue adaptation to proteolytic stress. Similarly, Rag1 KO mice lacking adaptive immune cells (*45*) displayed normal adaptation (Fig. 2C, S8A-B), indicating that lymphocytes are not essential for this process. Because proteases can signal via protease-activated receptor-2 (PAR2) (*46, 47*), we examined this pathway and found it dispensable for adaptation (Fig. S8C, D). Amphiregulin, an epidermal growth factor family member implicated in tissue repair (*48, 49*), was likewise not necessary (Fig. S8E-F). Given recent evidence that nociceptive neurons regulate tissue repair (*50–52*), we assessed their role using pharmacologic and genetic approaches. Resiniferatoxin (RTX) treatment effectively ablated nociceptive neuron function, confirmed by reduced capsaicin sensitivity and attenuated anaphylactic hypothermia (Fig. S9A), yet did not alter adaptation to proteolytic stress (Fig. S9B-C). Consistently, genetic ablation of TRPV1^+^ sensory neurons did not modify the acute proteolytic response (Fig. S9D-E). Collectively, these results exclude contributions of eosinophils, neutrophils, platelets, recruited monocytes, adaptive immune cells, PAR2 signaling, amphiregulin, and nociceptive neurons in mediating tissue adaptation to proteolytic stress.

**Fig. 2.**
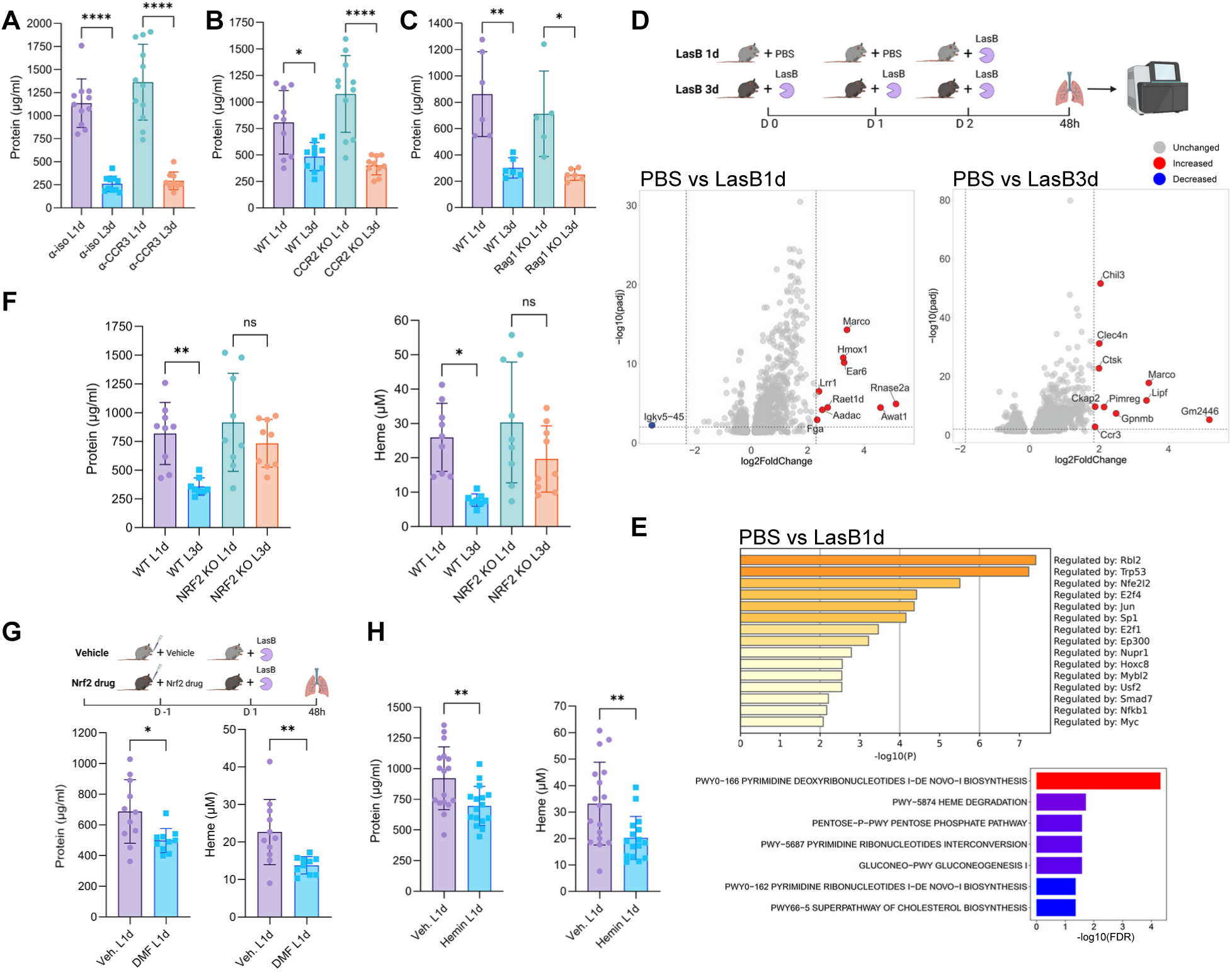
NRF2 is necessary for tissue adaptation to proteolytic stress. **(A-C)** BAL protein levels in animals treated with a-CCR3 antibody (A), in CCR2 (B) and Rag1 (C) KO mice after acute and repeated LasB exposure (n = 5-12 mice). **(D)** Experimental model of LasB tissue stress for bulk lung RNA-seq (D, top), volcano plots of differentially expressed genes after acute and repeated LasB exposure (D bottom) (n = 4 mice). **(E)** Transcription factor (top) and metabolic pathway (bottom) enrichment analyses of genes differentially expressed after acute LasB exposure. **(F)** BAL protein (left) and heme (right) levels in WT and NRF2 KO mice after acute and repeated LasB exposure (n = 8-9 mice). (G-H) Experimental model of pre-treatment with NRF2 agonists (G, top), BAL protein and heme levels with DMF (G bottom) and hemin (H) in animals pre-treated with DMF prior acute LasB exposure (n = 10-17 mice). Data are mean +/-SD and were analyzed by one-way ANOVA with Bonferroni multiple comparison test (A-C, F) and two-tailed unpaired t test (G-H). (*p=0.05, **p=0.01, ****p=0.0001).

To gain mechanistic insight, we performed bulk lung RNA-seq using our protocol of proteolytic tissue stress (Fig. 2D, top). Differential expression analysis revealed robust upregulation of heme oxygenase-1 (*Hmox1*) and macrophage-associated genes including *Marco*, *Ear6*, *Rnase2*, *Chil3*, and *Gpnmb* (Fig. 2D bottom). Transcription factor enrichment analysis using the TRRUST database (*53*) identified NRF2 as the top predicted regulator of stress-induced genes (Fig. 2E, top). NRF2 is a master antioxidant transcription factor activated during oxidative stress and previously implicated in lung responses to injury and inflammation (*54–58*). Gene ontology analysis revealed enrichment of pathways involved in heme degradation, consistent with elevated hemoglobin and heme levels (Fig. 1D-F), and the pentose phosphate pathway, which generates reducing equivalents for antioxidant defense (Fig. 2E bottom). To directly test the role of NRF2, we compared responses in WT and NRF2 KO mice. Strikingly, NRF2 KO animals failed to adapt to repeated proteolytic stress and maintained elevated BAL protein and heme levels (Fig. 2F). These mice also exhibited increased RBCs, hemoglobin, and hematocrit (Fig. S10A, C), along with sustained neutrophilia and depletion of AMs (Fig. S10B). To test activation of NRF2 pathway, we pre-treated animals with known NRF2 agonists (*56*), dimethyl fumarate (DMF) and hemin, before acute LasB exposure (Fig. 2G, top). Pre-treatment with NRF2 activators demonstrated protection from lung injury during acute proteolytic stress, as indicated by lower BAL protein and heme levels (Fig. 2 G, H). Similarly, pre-treated animals had lower levels of RBCs, platelets, hematocrit and hemoglobin as well as lower frequencies of neutrophils (Fig. S10 D-I). Together, these findings identify NRF2 as a central regulator of tissue adaptation to proteolytic stress.

### Alveolar macrophages mediate NRF2-dependent adaptation to proteolytic stress

To identify the cellular source of the NRF2-dependent transcriptional program, we performed single-cell RNA sequencing (scRNA-seq) after the protocol of proteolytic tissue stress. Unsupervised clustering based on canonical marker genes resolved epithelial, endothelial, mesenchymal, and immune compartments (Fig. 3A, S11). Among these populations, AMs emerged as the dominant cell type expressing NRF2 target genes, including canonical antioxidant and heme-detoxifying genes such as *Hmox1*, *Cat*, *Gclm*, *Blvra*, and *Blvrb* (Fig. 3B). Notably, we identified a proliferative subset of cycling AMs that accounted for the majority of the NRF2-dependent transcriptional response (Fig. 3B, S11C). Transcription factor enrichment analysis of differentially expressed genes in AMs confirmed NRF2 as the principal regulator of the transcriptional program (Fig. 3C, top). Gene ontology analysis further demonstrated enrichment of pathways related to heme degradation, glutathione biosynthesis, and the pentose phosphate pathway (Fig. 3C, bottom), consistent with NRF2-driven antioxidant metabolism and corroborating our bulk RNA-seq findings. To determine the necessity of AMs for tissue adaptation to proteolytic stress, we depleted AMs using clodronate liposomes, a well-established strategy for selective macrophage ablation (*59, 60*). Clodronate or control liposomes were administered prior to LasB exposure (Fig. 3D, top). At baseline, clodronate treatment caused a modest increase in BAL protein levels, consistent with congenital AM-deficient animal models (*61*), but did not affect RBC or platelet counts (Fig. S12A). Clodronate efficiently depleted AMs without substantially reducing other myeloid subsets (Fig. 3D). AM depletion resulted in a striking loss of adaptation: clodronate-treated animals exhibited markedly increased BAL protein and heme levels, along with elevated RBCs and platelets, during both acute and repeated proteolytic stress (Fig. 3E, S12B). These findings demonstrate that AMs are necessary for adaptation to proteolytic injury. Because commonly used Cre-driver lines targeting AMs (e.g., LysM^Cre^ and CD11c^Cre^) lack cell-type specificity and also affect neutrophils, monocytes, and dendritic cells (*62, 63*), we adopted an alternative strategy. We combined clodronate depletion with adoptive transfer of *ex vivo*-differentiated AMs to test the role of NRF2 during acute proteolytic tissue stress. Bone marrow cells from WT and NRF2 KO mice were differentiated into AMs and transferred into AM-depleted recipients, followed by LasB challenge the next day (Fig. 3F, top). Adoptive transfer restored AM numbers in BAL in both WT and NRF2 KO groups (Fig. S12E-F). This aligns well with prior studies showing that bone marrow-derived cells can replenish the AM niche and exhibit gene profiles, homeostatic, and phagocytic behavior similar to those of resident AMs (*64, 65*). However, transfer of NRF2 KO AMs failed to confer protection to the same extent as WT AMs, measured by cellular infiltration and BAL levels of hemoglobin, hematocrit, protein, and heme (Fig. 3F, S12C-D). To assess the relevance of this pathway in human disease, we interrogated publicly available transcriptomic datasets from patients with lung pathology (*66–68*). NRF2 target genes, HMOX1, BLVRA, and BLVRB, were consistently downregulated across multiple forms of human interstitial lung disease (ILD) (Fig. 3G). In fibrotic lungs, myeloid cells were the primary source of these transcripts (Fig. 3H). Moreover, HMOX1 expression was reduced in AMs from patients with COPD (Fig. 3I). Collectively, these findings establish alveolar macrophages as the principal cellular mediators of NRF2-dependent adaptation to proteolytic stress. NRF2 activity in AMs is necessary for vascular protection and tissue adaptation, and its dysregulation is evident across several human lung diseases characterized by impaired tissue repair.

**Fig. 3.**
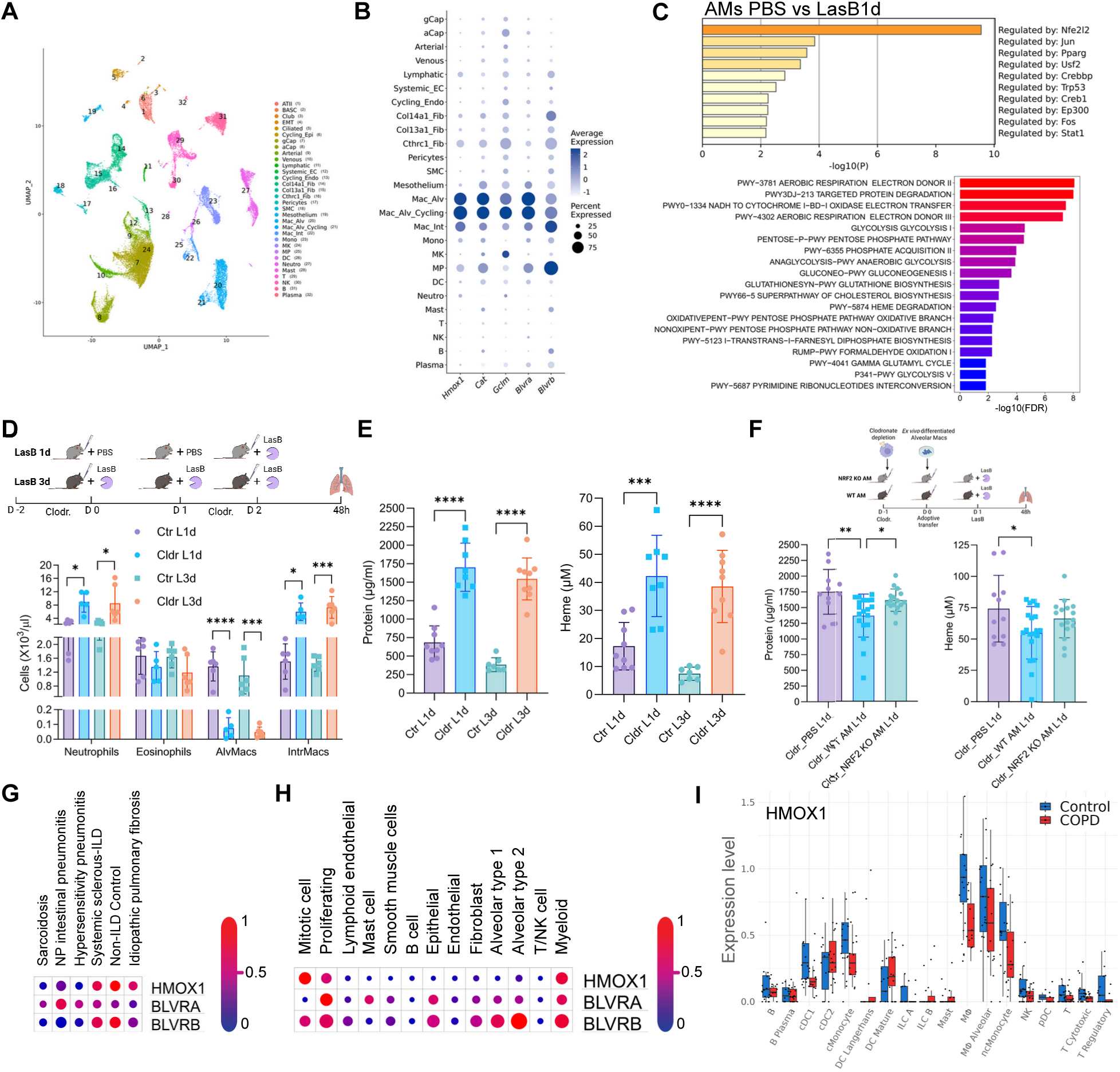
Alveolar macrophages are necessary for tissue adaptation to proteolytic stress. **(A-B)** UMAP of lung scRNA-seq showing annotated cell clusters (A) and dot plot showing expression of NRF2 target genes (B) after acute and repeated LasB exposure (n = 2-3 mice). **(C)** Transcription factor (top) and metabolic pathway (bottom) enrichment analyses of genes differentially expressed in AMs after acute LasB exposure. **(D-E)** Experimental model (D, top), lung myeloid cell numbers (D, bottom), BAL protein (E, left), and heme levels (E, right) following AM depletion and subsequent acute and repeated LasB exposure (n = 5-9 mice). **(F)** Experimental strategy (top), BAL protein (bottom left) and heme (bottom right) levels in animals after WT and NRF2 KO adoptive AM transfer and acute LasB exposure (n = 11-19 mice). **(G-H)** Dot plots showing expression of NRF2 target genes in human datasets across ILD diseases (*67*) (G) and cell populations (*66*) (H). (I) HMOX1 expression across immune cell populations in control and COPD patients (*68*). Data are mean +/- SD and were analyzed by one-way ANOVA with Bonferroni multiple comparison test (D-F). (*p=0.05, ***p=0.001, ****p=0.0001).

### NRF2 suppresses tissue-reparative signaling in macrophages

IL-4 stimulated macrophages play an important role in tissue repair (*69–71*). To test whether IL-4 – STAT6 signaling pathway plays a role in tissue protection to proteolytic damage, we compared responses to proteolytic stress in WT and STAT6 KO mice. Surprisingly, during acute proteolytic stress, STAT6 KO mice exhibited significantly lower BAL protein levels and reduced cellular infiltration, including RBCs, platelets, and neutrophils, relative to WT (Fig. 4A, S13A). Given that inflammatory pathways triggered by infectio us and sterile injury delay tissue repair programs until the source of the insult is eliminated (*70, 72*), our findings indicate that early suppression of STAT6-dependent reparative programs is beneficial during protease-induced oxidative stress. Since tuft cell-associated repair pathways have been implicated in injury responses across tissues (*73, 74*), we next examined the role of the tuft cell lineage-determining transcription factor Pou2f3 (*75*). Similar to STAT6 KO animals, Pou2f3 KO mice displayed reduced BAL protein, leukocyte infiltration, RBCs, and platelets during acute proteolytic stress (Fig. 4B, S13B). These results further support the concept that tissue-repair programs are incompatible with the acute oxidative environment induced by proteolytic injury.

**Fig. 4.**
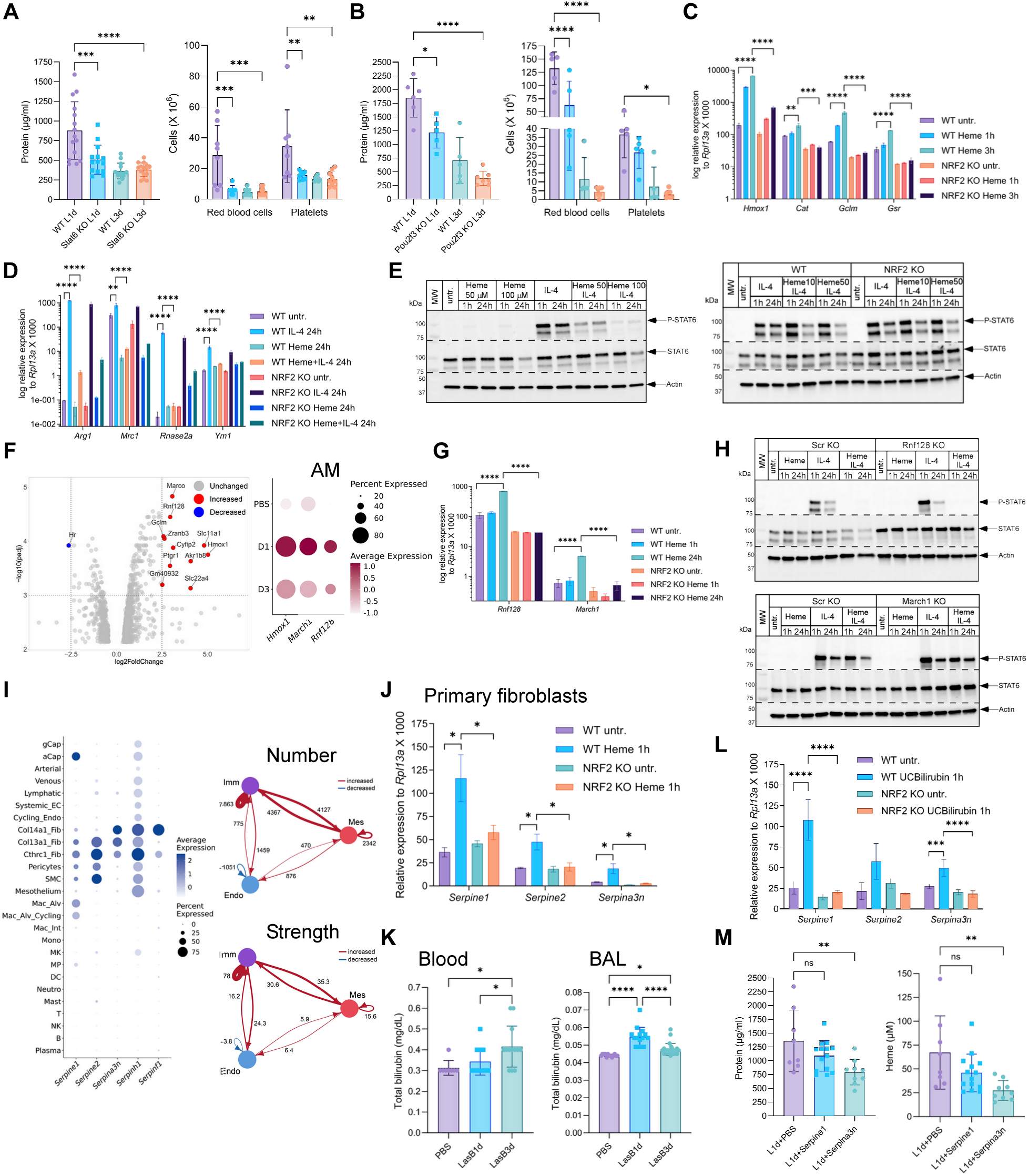
NRF2 regulates tissue-reparative program in macrophages and anti-proteolytic response in fibroblasts. **(A-B)** BAL protein levels and RBC and platelet counts in WT and STAT6 (A) and Pou2f3 (B) KO mice after acute and repeated LasB exposure (n = 5-15 mice). **(C)** Relative mRNA levels of NRF2 target genes in WT and NRF2 KO BMDMs stimulated with heme (100 µM) for indicated times. **(D)** Relative mRNA levels of STAT6 target genes in WT and NRF2 KO BMDMs stimulated with heme (100 µM), IL-4 (20 ng/ml) and both for 24 h. **(E)** WB of total and P-STAT6 levels in WT (left) and NRF2 KO (right) BMDMs stimulated with heme, IL-4 (20 ng/ml) and both for indicated times. **(F)** Volcano plot of differentially expressed genes (left) and dot plot showing expression of *Hmox1*, *Rnf128* and *March1* (right) in AMs following LasB exposure. **(G)** Relative mRNA levels of *Rnf128* and *March1* in WT and NRF2 KO BMDMs stimulated with heme (100 µM) for indicated times. **(H)** WB of total and P-STAT6 protein levels in Scr and Rnf128 (top) and March1 (bottom) KO BMDMs stimulated with heme (100 µM), IL-4 (20 ng/ml) and both for indicated times. **(I)** Dot plot of lung scRNA-seq showing expression of *Serpin* genes (left) after repeated LasB exposure. The inferred number (top right) and strength (bottom right) of signaling change between immune (purple circle), mesenchymal (red circle), and endothelial (blue circle) cells after acute LasB exposure, quantified by Cell Chat. Edge width denotes the relative number and interaction strength. **(J)** Relative mRNA levels of *Serpin* genes in WT and NRF2 KO primary lung fibroblasts stimulated with heme (100 µM) for 1 hr. **(K)** Total bilirubin levels in blood (left) and BAL (right) after acute and repeated LasB exposure (n = 8-24 mice). **(L)** Relative mRNA levels of *Serpin* genes in WT and NRF2 KO primary lung fibroblasts stimulated with unconjugated bilirubin (10 µM) for 1 hr. (M) BAL protein (left) and heme (right) levels in mice treated with Serpine1 or Serpina3n after LasB exposure (n = 8-14 mice). mRNA expression was measured relative to *Rpl13a*. ß-actin served as a loading control. Data are mean +/- SD and were analyzed by one-way ANOVA with Bonferroni multiple comparison test. (*p=0.05, **p=0.01, ***p=0.001, ****p=0.0001).

To define the molecular basis of this suppression, we modeled oxidative stress *in vitro* using bone marrow–derived macrophages (BMDMs) that were stimulated with NRF2 agonists (DMF, heme), IL-4, or both. Heme or DMF induced a robust NRF2 -dependent transcriptional program, including upregulation of *Hmox1*, *Cat*, *Gclm*, and *Gsr* (Fig. 4C, S13C). Notably, NRF2 activation by heme or DMF suppressed IL-4-induced STAT6 target genes (*Arg1*, *Mrc1*, *Rnase2a*, *Ym1*), and this inhibition was partially relieved in NRF2-deficient BMDMs (Fig. 4D, S13D). This inhibition was accompanied by reduced phosphorylation of STAT6 (P-STAT6) at an early time point (1hr) and decreased total STAT6 protein levels at 24hr. In contrast, both phosphorylated and total STAT6 were more stable in NRF2 KO BMDMs (Fig. 4E), suggesting that NRF2 promotes STAT6 inactivation and degradation. Because STAT6 regulation can occur through phosphatase or ubiquitin-mediated degradation pathways, we examined our scRNA-seq dataset and identified induction of the ubiquitin ligase genes *Rnf128* and *March1* following LasB exposure in AMs (Fig. 4F). Expression of both these genes was dependent on NRF2 in BMDMs (Fig. 4G). Previous studies have shown that Rnf128 promotes STAT6 degradation in Th2 cells, whereas March1 downregulates cell surface immune receptors via lysosomal pathways (*76, 77*). To directly test whether these ligases regulate STAT6 stability in macrophages, we generated Rnf128, March1, and control (scramble-Scr) KO macrophages differentiated from Cas9-expressing ER-Hoxb8 myeloid progenitors (*78*), and stimulated them with heme, IL-4, or both. By measuring P-STAT6 and total STAT6 levels, we found that these two ubiquitin ligases targeted different forms of STAT6 for degradation. Rnf128 KO BMDMs showed increased stabilization of total STAT6 levels (Fig. 4H, top), whereas March1 KO had a stronger effect on stabilization of P-STAT6 (Fig. 4H bottom). These findings suggest that Rnf128 primarily targets total STAT6 for degradation, whereas March1 preferentially targets the phosphorylated form. Consistent with this, both knockouts enhanced IL-4-induced STAT6 target gene expression, and March1-deficient BMDMs were partially protected from heme-mediated suppression of IL-4 responses (Fig. S13E-F) Collectively, these data demonstrate that NRF2 antagonizes IL-4-STAT6-dependent reparative signaling in macrophages through induction of the ubiquitin ligases.

### NRF2 regulates anti-proteolytic response in fibroblasts

To determine how proteolytic activity is restrained during tissue adaptation, we profiled protease inhibitor expression in our scRNA-seq and bulk RNA-seq datasets. We identified induction of multiple Serpin family members, including *Serpine1*, *Serpine2*, *Serpina3n*, *Serpinh1*, and *Serpinf1* (Fig. 3I, S15A). Serpins are canonical endogenous protease inhibitors and therefore represent promising candidates to prevent proteolytic stress. Among these candidates, Serpinh1 and Serpinf1 encode non-inhibitory serpins involved in collagen folding and angiogenesis, respectively. Serpine1 and Serpine2 are closely related regulators of fibrinolysis and clot formation, whereas Serpina3n encodes a secreted inhibitor with broad protease specificity (*79, 80*). scRNA-seq revealed that mesenchymal cells were the principal source of *Serpin* induction during proteolytic stress (Fig. 3I, S14), with distinct subsets of alveolar fibroblasts exhibiting the highest expression levels (Fig. S15B-C). Notably, interrogation of publicly available human dataset (*67*) demonstrated reduced *Serpin* expression in ILD samples (Fig. S15D), suggesting impaired protease inhibitory capacity in human lung pathology.

AMs expressed high levels of NRF2-dependent heme-catabolic genes (*Hmox1*, *Blvra*, *Blvrb*), whereas subsets of alveolar fibroblasts expressed these genes at comparatively low levels (Fig. S14C), implying functional specialization between these cellular compartments. Consistent with this, scRNA-seq-based cell-cell communication analysis revealed enhanced immune-mesenchymal signaling during repeated proteolytic stress (Fig. 3I), and spatial analysis demonstrated closer association between macrophages and fibroblasts following LasB exposure (Fig. S17). To test whether fibroblasts directly respond to heme and to define the mechanism of Serpin induction, we isolated primary lung fibroblasts from WT and NRF2 KO mice and stimulated them with heme, alongside BMDMs. Heme robustly induced *Serpin* expression in fibroblasts in an NRF2-dependent manner (Fig. 4J). In contrast, induction of canonical heme-catabolic genes was modest (*Hmox1*) or absent (*Blvrb*) in fibroblasts, whereas BMDMs strongly upregulated *Hmox1* and *Blvrb* but did not induce *Serpins* (Fig. S16A–B). The NRF2-driven heme catabolic pathway converts heme into Fe²^+^, CO, and biliverdin via Hmox1, followed by reduction of biliverdin to bilirubin by Blvra/Blvrb (*81*). These data suggest a model in which macrophages primarily detoxify heme and generate bilirubin, thereby limiting heme -mediated cytotoxicity in neighboring fibroblasts. Consistent with this, AM depletion resulted in significantly elevated heme levels in the BAL, indicating impaired heme clearance in the absence of AMs (Fig. 3E). Consistent with enhanced heme catabolism, bilirubin concentrations increased in blood and BAL during proteolytic stress (Fig. 4K). Strikingly, fibroblasts responded to unconjugated (hydrophobic) bilirubin but not soluble conjugated bilirubin by inducing *Serpin* expression in an NRF2-dependent manner (Fig. 4L, S16C), suggesting that macrophage-derived bilirubin acts as a signal to promote fibroblast-mediated *Serpin* expression.

To assess whether candidate Serpins directly inhibit LasB activity, we performed elastase cleavage assays *in vitro* in the presence of recombinant Serpins and phosphoramidon, a known LasB inhibitor. Phosphoramidon potently suppressed LasB activity, whereas Serpine1 exhibited no detectable inhibition and Serpina3n demonstrated intermediate inhibitory capacity (Fig. S16D). *In vitro* cleavage assays revealed that Serpine1 was fully degraded by LasB, whereas Serpina3n underwent specific cleavage with formation of higher-molecular-weight complexes (Fig. S16E), consistent with covalent complex formation between the Serpina3n reactive center loop and the LasB catalytic site. Complex formation was dose dependent (Fig. S16F), confirming that Serpina3n functions as a LasB inhibitor (*82*). To test the protective capacity of these Serpins *in vivo*, we administered recombinant Serpins following LasB exposure. Serpina3n-treated animals exhibited markedly reduced BAL protein, heme, hemoglobin, hematocrit, RBCs, and platelet levels compared to PBS-treated controls, whereas Serpine1 conferred partial protection (Fig. 4M, S16G-H). Serpina3n treatment also reduced neutrophil frequencies and increased AM percentage (Fig. S16I). Collectively, these findings establish a coordinated protective axis in which NRF2 activation in macrophages drives heme detoxification and bilirubin production, whereas NRF2 activation in fibroblasts induces Serpin expression and most prominently Serpina3n to directly inhibit protease activity.

### Tissue adaptation to proteolytic stress confers broad protection against subsequent injury and infection

To determine the functional consequences of adaptation to proteolytic tissue stress, we subjected mice to secondary challenges one week after completion of the proteolytic stress protocol. We first evaluated susceptibility to an increased acute dose of LasB (0.1 mg/kg) (Fig. 5A, top). Mice adapted to proteolytic stress (LasB3d) were markedly protected from vascular injury compared to LasB1d and PBS groups. Adapted animals exhibited significantly lower BAL protein, erythrocyte, platelet, hemoglobin, hematocrit, and heme levels following high -dose LasB challenge (Fig. 5A-C). This protection was accompanied by reduced neutrophil infiltration and increased frequencies of AMs (Fig. 5D, S18A), consistent with the protective macrophage phenotype during adaptation. We next assessed response against *Pseudomonas aeruginosa* (PA01 strain) infection (Fig. 5E). Similar to the high-dose protease challenge, adapted LasB3d animals demonstrated enhanced protection, characterized by reduced bacterial burden in BAL and lung tissue, as well as lower BAL protein and erythrocyte levels ( Fig. 5E-G). These differences were not associated with changes in body temperature, total leukocyte infiltration, or neutrophil frequencies, however, adapted animals maintained increased AM representation (Fig. S18B-C). Finally, we examined susceptibility to viral infection using influenza A virus (WSN strain) (Fig. 5I). Adapted LasB3d animals experienced reduced body weight loss and hypothermia and exhibited improved recovery compared to controls (Fig. 5I). Survival was also enhanced (Fig. S18D). Similar protection was observed with a second influenza strain (PR8) (Fig. S18E). To determine the requirement for NRF2 in this protective response, we compared adapted LasB3d WT and NRF2 KO mice following influenza infection. NRF2 KO animals failed to maintain protection and exhibited greater weight loss, hypothermia, and reduced survival (Fig. 5J–K, S18F). These findings demonstrate that NRF2 is necessary for the sustained protective state induced by prior proteolytic stress. Collectively, these results show that adaptation to proteolytic tissue stress establishes a durable protective program, consistent with prior studies of inflammatory injury(*20, 21, 83*), that confers cross-protection against subsequent protease challenge as well as bacterial and viral infections.

**Fig. 5.**
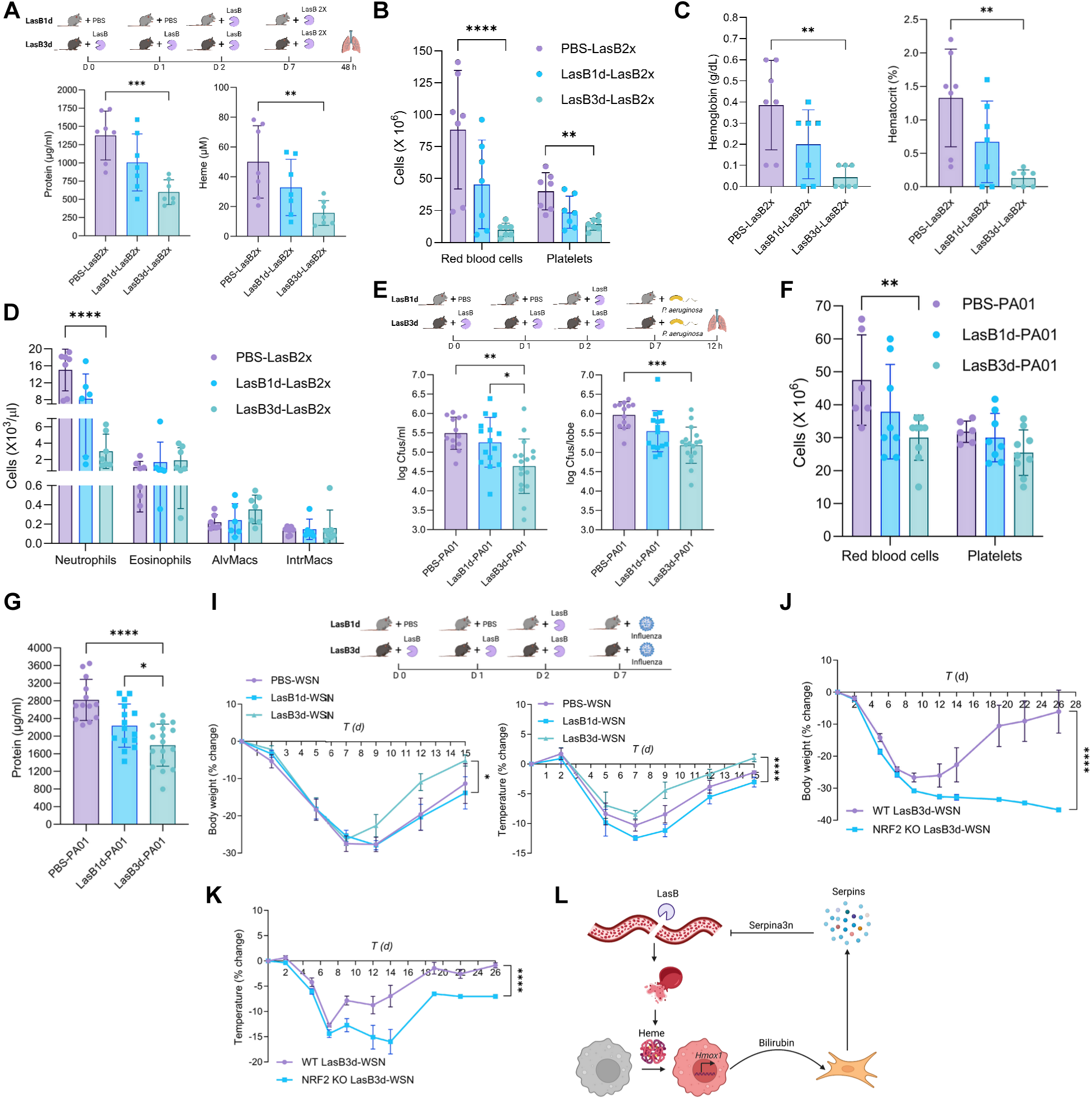
Tissue adaptation to proteolytic stress provides protection against subsequent challenges. **(A-D)** Experimental model of subsequent acute high-dose protease exposure (A, top), BAL protein (A, bottom left) and heme levels (A, bottom right), RBC and platelet counts (B), hemoglobin and hematocrit levels (C), and myeloid cell numbers (D) in animals challenged with high-dose LasB 1 week after prior acute or repeated exposure (n = 6-7 mice). **(E-G)** Experimental model of subsequent acute *P. aeruginosa* infection (E, top), BAL (E, bottom left) and lung (E, bottom right) bacterial load, BAL RBC and platelet counts (F) and protein levels (G) in animals challenged with *P. aeruginosa* infection 1 week after prior acute or repeated LasB exposure (n = 8-17 mice). **(I)** Experimental model of subsequent inlfuenza infection (I, top), kinetics of body weight change (I, bottom left) and temperature change (I, bottom right) in animals challenged with WSN influenza infection 1 week after prior acute or repeated LasB exposure (n = 7-8 mice). **(J-K)** Kinetics of body weight change (J) and temperature change (K) in WT and NRF2 KO animals challenged with WSN influenza infection 1 week after prior repeated LasB exposure (n = 6 mice). **(L)** Model of tissue adaptation to proteolytic stress. Data are mean +/- SD and were analyzed by one-way ANOVA with Bonferroni multiple comparison test (A-G) and two-way ANOVA (I-K). (*p=0.05, **p=0.01, ***p=0.001, ****p=0.0001).

## Discussion

In this study, we identify vascular disruption and extracellular heme release as a hallmark of acute proteolytic stress in the lung. Rather than representing irreversible damage, this injury initiates an adaptive program upon repeated exposure. We delineate the cellular and molecular basis of this adaptation as an intercellular stress-response circuit centered on AMs and fibroblasts (Fig. 5L). Vascular injury releases heme, which is sensed by AMs and triggers an NRF2 - dependent oxidative stress response. Activation of NRF2 induces *Hmox1* and *Blvra*/*Blvrb*, enabling conversion of cytotoxic heme into cytoprotective metabolite bilirubin. Thus, proteolytic tissue damage is translated into a metabolically coupled antioxidant response. Bilirubin acts on neighboring fibroblasts and induces *Serpin* expression. Among these, Serpina3n emerges as the most potent inhibitor of proteolytic activity and the most effective in limiting tissue injury *in vivo*. Together, these findings reveal a cooperative protective network where macrophages sense and detoxify heme, generating a secondary signal, and fibroblasts execute the effector arm by producing Serpins that suppress proteolytic damage.

Our model extends classical observations that protease exposure disrupts basement membrane and causes vascular injury and hemorrhage (*16, 18*). Our findings suggest that vascular injury and heme release may represent a generalizable feature of proteolytic stress across tissues, as protease-induced hemorrhage has been reported in skin and intestine in addition to the lung (*84, 85*). Comparative analysis will be necessary to define shared versus organ-specific patterns of proteolytic stress adaptation.

Protease-antiprotease imbalance is a defining feature of diverse lung pathologies, including viral infections, acute respiratory distress syndrome (ARDS), alveolar hemorrhage, COPD, and ILD (*30, 31*). We demonstrate that protease exposure activates an NRF2-driven oxidative stress program, consistent with elevated oxidative stress signatures observed in ARDS, COPD, and ILD (*86–88*). Thus, our studies provide a mechanistic bridge between proteolytic injury and NRF2 activation. Analysis of publicly available human lung disease datasets (*66–68*) further supports enrichment of NRF2 target genes in myeloid cells, aligning with our finding that AMs are the principal source of the NRF2 gene program. Although AM numbers transiently decline following acute protease exposure, similar to viral, bacterial, or bleomycin injuries (*33, 35, 36, 38*), the reduction is comparatively modest and does not depend on infiltrating monocytes or monocyte-derived macrophages. Instead, we detect a population of cycling AMs and maintenance of macrophage numbers upon repeated challenge, suggesting local proliferation as a mechanism to restore AM numbers and function during repeated stress. This observation is consistent with previous models of AM replenishment that depends on the degree of initial depletion (*35, 37, 38, 89, 90*).

Our data further highlight a functional asymmetry between macrophages and fibroblasts in heme handling. Our findings establish that AMs possess the complete enzymatic machinery for heme catabolism and are indispensable for limiting extracellular heme accumulation in the lung. Macrophage depletion leads to marked increases in tissue heme burden, corroborating extensive prior evidence that macrophages are essential for heme clearance (*81, 91–93*). Although fibroblasts can induce *Hmox1*, they lack the full enzymatic machinery to metabolize heme efficiently (*94, 95*). Instead, fibroblasts rely on macrophages to detoxify heme and convert it to bilirubin which induces *Serpin* expression in NRF2-depdendet manner. Notably, NRF2 activation generates distinct transcriptional outputs in macrophages and fibroblasts, underscoring cell-type-specificity of a shared stress pathway. This division of labor ensures coordinated detoxification and protease inhibition.

Among fibroblast populations, alveolar fibroblasts and Cthrc1^+^ fibroblasts are the principal sources of Serpins. Although Cthrc1^+^ fibroblasts have been implicated in fibrotic remodeling and adverse outcomes (*96*), our findings suggest that during early injury they may exert protective effects by supplying antiproteases that restore tissue homeostasis. Dysregulation of this protective axis at later stages could contribute to pathological remodeling, highlighting context-dependent roles for fibroblast subsets during injury and repair.

The exact molecular mechanism by which bilirubin induces *Serpin* expression remains unresolved. Several candidate bilirubin receptors have been proposed (*97, 98*), but their relative contributions in lung fibroblasts require further investigation. Among fibroblast-derived Serpins, Serpina3n is particularly effective in inhibiting proteolytic activity and limiting damage *in vivo*. Recent studies indicate additional protective roles for Serpina3n in enhancement of the blood-brain barrier and attenuation of neuropathic pain (*99, 100*). The specificity of Serpina3n toward diverse protease classes and its regulation during infectious and sterile inflammation warrant future study. These questions may inform therapeutic strategies that combine NRF2 agonists, already in clinical use across diverse disease contexts, with Serpin supplementation to reinforce tissue protection.

Finally, adaptation to proteolytic stress confers broader protection against subsequent high -dose protease as well as viral and bacterial infections, indicating convergence of diverse injuries on a shared NRF2-dependent program (*30, 54, 57, 86–88, 93*). Defining the tissue-level mechanisms underlying this cross-protection will clarify whether NRF2-driven circuits represent a general principle of injury adaptation across inflammatory contexts and protease classes.

In summary, we define vascular damage and heme release as initiating events in proteolytic stress and delineate a lung-specific intercellular circuit in which AMs sense and detoxify heme and fibroblasts produce Serpin protease inhibitors to suppress proteolytic injury. This cooperative macrophage–fibroblast axis provides a mechanistic framework for understanding how tissues detect and adapt to proteolytic damage and offers potential avenues for therapeutic modulation of stress adaptation pathways across inflammatory diseases.

## Acknowledgments

We are grateful to all members of the Medzhitov laboratory for thoughtful discussions; E. Condiff, S. Erickson, E. Kopp, I. Licona-Limon, S. Ovadia, D. Ryan, B. Vaidyanathan for useful suggestions and help with preliminary experiments; Y. Yuan for initial assistance with analysis of scRNA-seq; Cuiling Zhang, Chuck and Sophie Annicelli, and Jaime Cullen for mouse colonies and help with *in vivo* experiments. We thank Dr. B. Kazmierczak (Yale University) for providing *P. aeruginosa* genomic DNA, Dr. C. Wilen (Yale University) for Pou2f3 KO mice and Dr. A. Iwasaki (Yale University) and members of the Iwasaki laboratory for help in using Piccolo Express analyzer. We thank Dr. D. Zaiss (University of Regensburg) for the Areg KO mice. We also thank Yale Flow Cytometry for their assistance with FACS service and cell sorting, Yale Pathology Tissue Services for their assistance with histological analysis, the Yale Translational Research Imaging Center for their assistance with µCT, and the Keck Microarray Shared Resource (KMSR) at Yale University and the Yale Center for Genome Analysis (YCGA) for their assistance with bulk and scRNA-seq services. Images were created with BioRender.com.

## Funding

The Howard Hughes Medical Institute (RM)

The Blavatnik Family Foundation (RM)

The Food Allergy Science Initiative (RM)

National Institutes of Health grant R01AI144152 (RM)

National Institutes of Health grant R01AI182061 (RM)

National Institutes of Health grant P01AI179273 (RM)

Yale Brown-Coxe Post-Doctoral Fellowship (KA)

Howard Hughes Medical Institute Fellowship of Life Sciences Research Foundation (KA)

National Institutes of Health grant HL126094 (CSDC)

The Department of Defense Grant (CSDC)

Francis B Parker Fellowship (LS)

American Lung Association Catalyst Award (LS)

## Author contributions

Conceptualization: KA, RM

Methodology: KA, AMG, SY, ARC, DWEN, EPM, LS

Investigation: KA, AMG, SY, ARC, DWEN, EPM, LS

Visualization: KA, AMG, EPM

Funding acquisition: RM, KA, CSDC, LS, EPM

Project administration: RM, KA

Supervision: RM

Writing – original draft: KA, RM

Writing – review & editing: KA, RM, AMG, EPM

## Data, code, and materials availability

All data are available in the main text or the supplementary materials. Bulk sequencing and scRNA-seq of lung generated for this paper can be accessed through the Gene Expression Omnibus at accession number. All original code is publicly available on Github.

## Supplementary Materials

Materials and Methods

Supplementary Text

Figs. S1 to S18

Table S1

## Materials and Methods

### Animals and animal models

Mice were bred at the Yale Animal Resources Center at Yale University in specific pathogen-free conditions, and all experiments were done in accordance with approved guidelines, regulations, and protocols as determined by the Institutional Animal Care and Use Committee at Yale. C57BL/6 and BALB/cJ mice were bred in-house. NRF2^-/-^ (strain# 017009), CCR2^-/-^ (strain# 004999), Rag1^-/-^ (strain# 002216), ΔdblGATA (strain# 033551), PAR2^-/-^ (strain# 004993), Stat6^-/-^ (strain# 005977), TRPV1^Cre^ (strain# 017769), ROSA^DTA^ (strain# 009669) were purchased from Jackson Laboratories. Areg^-/-^ C57BL/6 mice were kindly provided by Dr. Zaiss (University of Edinburgh). Pou2f3^-/-^ C57BL/6 mice were kindly provided by Dr. Wilen (Yale University).WT animals were either in-house bred C57BL/6 or BALB/cJ mice and/or littermate wild type/heterozygotes. Mice 8-12 weeks of age were used for all experiments; all animals were age- and sex-matched and then randomized into the different groups.

#### Model of proteolytic tissue stress

Mice were anesthetized with a ketamine (100mg/kg)/xylazine (10mg/kg) mixture and 40 µl of LasB (0.05 mg/kg), subtilisin (0.25 mg/kg; Sigma-Aldrich) or papain (2 mg/kg; Thermo Fisher Scientific) was administered dropwise intranasally once (acute stress) or three times (repeated stress) within a 72-hour interval.

#### Pre-treatment with NRF2 agonists

Mice were anesthetized with a ketamine (100mg/kg)/xylazine (10mg/kg) mixture and DMF (5 mg/kg; Cayman Chemical) or hemin (55 µg/kg; Frontier Specialty Chemicals) and vehicle controls were administered dropwise intranasally. 24 h later mice were anesthetized as above and 40 µl of LasB (0.05 mg/kg) was administered dropwise intranasally. DMSO served as vehicle control for DMF and NaOH:HCl (4:3) – for hemin.

#### Depletion of alveolar macrophages

Mice were anesthetized with a ketamine (100mg/kg)/xylazine (10mg/kg) mixture and 40 µl of control or clodronate liposomes (11 mg/kg, Encapsula Nanosciences CLD-8909) were administered dropwise intranasally. 24 h later mice were anesthetized as above and 40 µl of LasB (0.05 mg/kg) was administered dropwise intranasally. For the repeated LasB stress, depletion was performed prior to the first and the last LasB administrations.

#### Adoptive transfer of alveolar macrophages

Mice were anesthetized with a ketamine (100mg/kg)/xylazine (10mg/kg) mixture and 40 µl of clodronate liposomes (Encapsula Nanosciences CLD-8909) was administered dropwise intranasally. 24 h later mice were anesthetized as above and 40 µl of *ex vivo*-differentiated alveolar macrophages (AMs) (1-x10^6^) was administered dropwise intranasally.

#### Serpin-mediated inhibition of proteases in the model of acute proteolytic stress

Mice were anesthetized with a ketamine (100mg/kg)/xylazine (10mg/kg) mixture and 40 µl of LasB (0.05 mg/kg) was administered dropwise intranasally. 24 h later mice were anesthetized as above and 40 µl of PBS, Serpine1 (0.5 mg/kg) or SerpinA3n (1 mg/kg) was administered dropwise intranasally.

#### Antibody-mediated depletion of eosinophils and neutrophils

a-CCR3 and corresponding isotype control antibodies (15 mg/kg; BioXCell), as well as a-Ly6G and corresponding isotype control antibodies (11 mg/kg; BioXCell), were administered intraperitoneally. 24 h later mice were anesthetized with ketamine (100 mg/kg)/xylazine (10 mg/kg), and 40 µl LasB (0.05 mg/kg) was administered intranasally dropwise. For repeated LasB stress, depletion was performed prior to the first and the last LasB administrations.

#### Clopidogrel treatment

Clopidogrel (5 mg/kg; Cayman Chemical) or vehicle control (DMSO:PBS – 1:2) were administered intraperitoneally. 24 h later mice were anesthetized with ketamine (100 mg/kg)/xylazine (10 mg/kg), and 40 µl LasB (0.05 mg/kg) was administered intranasally dropwise. For repeated LasB stress, depletion was performed prior to the first and the last LasB administrations.

#### Resinferatoxin-mediated depletion of nociceptive neurons

Resiniferatoxin or vehicle control (AdipoGen Life Sciences) were administered to 4-week-old mice on three consecutive days (30, 70, and 100 µg/kg in 100 µl PBS with 2% DMSO and 0.15% Tween 80, flank injection). Four weeks later, mice were subjected to the LasB tissue stress protocol as above.

#### Eye-wipe test

10 µl of capsaicin (Cayman Chemical) solution or vehicle (300 µM in PBS) were applied to the mouse cornea, and eye wipes were counted over 30 s.

#### Capsaisin-mediated anaphylaxis

Capsaicin (2 mg/kg in 100 µl PBS with 2% DMSO and 0.15% Tween 80; Cayman Chemical was administered intraperitoneally. Rectal temperatures were followed for 0.5-1 h after challenge.

#### Acute high-dose LasB challenge

7 days after acute and repeated LasB stress protocol, animals were anesthetized as above and 40 µl of LasB (0.1 mg/kg) was administered dropwise intranasally.

#### PAO1 infections

7 days after acute and repeated LasB stress protocol, mice were infected with 2.5 x 10^6^ CFUs of bacteria per mouse in 50 ul of PBS by intratracheal route. Briefly, mice were anesthetized using ketamine and xylazine (100 and 10 mg/kg, respectively). A vertical cut on the neck was made after ensuring proper anesthesia by toe pinch to exposure the trachea. 50 µl of PBS containing bacteria were instilled directly in the trachea using a Hamilton syringe. The cut was sealed suing 3M Vetbond glue (3M). Mice were euthanized at 12 hours post infection to harvest BAL and lung samples to enumerate number of bacteria and inflammation.

#### Influenza infections

7 days after acute and repeated LasB stress protocol, mice were anesthetized as above and infected with a sublethal dose of 300 PFU of A/WSN/33 (H1N1) strain of influenza A virus in a volume of 30 µl intranasally. For PR8 strain (H1N1), mice were anesthetized using ketamine/xylazine solution as above and an infection inoculum of 10 PFUs in 50 µl of PBS was administered by intranasal route. Changes of body weight and temperature were recorded over the course of infections.

### Protein expression and purification

#### LasB expression

The coding sequence of *P. aeruginosa* ( was amplified by PCR using forward (CTGCTAGCAAGAAGGTTTCTACGCTTGACCTG) and reverse (GAAAGCTTCAACGCGCTCGGGCAGG) primers containing NcoI and HindIII restriction digestion sites, respectively, from *P. aeruginosa* PAO1 genomic DNA and cloned into the expression vector pET21d (Novagen) to express C-terminally His6-tagged version of a protein. LasB was expressed using T7 Express LysY (New England Biolabs) or BL21 (DE3) pLysS (Promega) cells. The overnight cell culture (2-5 ml) was used to inoculate 0.2-0.8L of LB and the cell culture was incubated for 3-4 hr at 37 °C until OD at 600 nm reached 0.6-0.8 units. The protein expression was then induced by addition of 0.2 mM IPTG (Sigma-Aldrich or Cayman Chemical). The cells were harvested after 4 hr of incubation at 37 °C and either kept at -20 °C or disrupted by sonication. LasB was first purified by affinity chromatography on Ni^2+^-agarose beads (Qiagen) followed by anion exchange chromatography in 50 -500 mM gradient of NaCl on Mono Q column (GE Healthcare). The eluted fractions were concentrated to 1-3 µg/µl concentration and endotoxin was removed using Pierce High Capacity Endotoxin Removal Resin (Thermo Fisher Scientific). After endotoxin removal, LasB was aliquoted and stored at -80 °C until use. Endotoxin levels were measured using Pierce LAL Chromogenic Endotoxin Quantitation Kit (Thermo Fisher Scientific).

#### Serpine1 and Serpina3n expression

cDNA for Sepine1 and Serpina3n with C-terminal 6XHis-tag (Twist Biosciences) was cloned into pcDNA3.1 vector containing CMV promoter. The plasmids were transfected into Expi293 cells according to manufacturer’s instructions (Thermo Fisher Scientific). For protein purification, Expi293 cells expressing recombinant His-tagged proteins were collected at day 4-6 post-transfection and pelleted by centrifugation (5 min, 500 × g, 4 °C). The clarified supernatant was passed through a 0.22 µm filter (Corning) to remove residual cellular debris. The supernatant was supplemented with imidazole (final concentration 10 mM) and NaCl (10 mM). Ni²^+^-charged agarose beads (Qiagen) were prepared by washing twice with 50 ml Milli-Q water followed by 25–50 ml low-salt wash buffer (25 mM Tris, pH=8.0; 300 mM NaCl, 15 mM imidazole) in 50 ml conical tubes. Approximately 1 ml of settled Ni-agarose resin was used per 300 ml of filtered supernatant. The prepared supernatant was added to the equilibrated Ni-agarose beads and incubated with gentle mixing for 3–4 h at 4 °C to allow binding of His-tagged proteins. All subsequent steps were performed in a cold room (4 °C). The bead–protein mixture was applied to the column and washed with 25 ml Ni low-salt wash buffer (=10 column volumes). The bound proteins were eluted in 0.5 ml fractions using elution buffer (25 mM Tris, pH=8.0; 300 mM NaCl, 0.25 M imidazole). Protein-containing fractions were monitored by measuring absorbance at 280 nm. Protein purity was assessed by SDS–PAGE under denaturing conditions.

### Cell culture

#### Bone marrow-derived macrophages

Bone marrow was isolated from the tibia and femur of 8-12 week-old male and female mice. Bone marrow cells were isolated by flushing and plated for macrophage differentiation in non–tissue culture-treated Petri dishes and cultured for 7 days in RPMI (Corning) supplemented with 10% fetal bovine serum (Gemin Bio), 30% L929 conditioned medium, L-glutamine, penicillin/streptomycin, non-essential amino acids, HEPES, and sodium pyruvate (all Gibco) to generate bone marrow-derived macrophages (BMDMs). On day 7, macrophages were harvested by removing culture medium and incubating cells with ice - cold PBS containing 5 mM EDTA for 3-5 min. Cells were then replated and used for downstream applications the following day.

#### Bone marrow-derived alveolar macrophages

Alveolar macrophages were generated from bone marrow in the same RPMI medium supplemented with GM-CSF (20 ng/ml, Genescript) and TGF-ß (2 ng/ml, Genescript). On day 7, medium was refreshed with GM-CSF and TGF-ß and supplemented with the PPARγ ag onist rosiglitazone (0.1 µM, Cayman Chemical) for an additional 2 days. Cells were subsequently passaged every 3 -4 days in medium containing GM-CSF, TGF-ß, and rosiglitazone and maintained for 2-4 months.

#### ER-HoxB8 myeloid progenitors

Cas9-expressing ER-HoxB8 were maintained in progenitor medium consisting of RPMI (Corning) supplemented with 10% fetal bovine serum (GeminiBio), 2 mM L-glutamine, 1 mM sodium pyruvate, 10 mM HEPES (all Gibco), ß-mercaptoethanol (Sigma-Aldrich), 2% GM-CSF-conditioned medium, and 2 µM ß-estradiol (Sigma-Aldrich). Myeloid progenitors were differentiated into macrophages according to the protocol for BMDMs (see above). To maintain optimal differentiation efficiency, progenitor cells were kept at densities below 0.5 × 106 cells/ml prior to plating, with minimal debris and aggregation. Suspension cells were collected, excluding adherent cells, and washed twice in PBS containing 1% fetal bovine serum. Cells (3 × 106) were plated in 15 cm non-tissue culture–treated dishes in 20 ml macrophage differentiation medium as above. Cells were differentiated for 7–9 days, with an additional medium supplementation on day 3 and 6 when cultures were maintained beyond day 7. For harvest, non-adherent cells were first removed by washing with warm PBS, followed by detachment of adherent cells using trypsin/EDTA at 37°C for approximately 5 min.

*Primary lung fibroblasts* were isolated from mouse lungs following enzymatic digestion. Briefly, lungs were perfused with cold PBS, excised, minced into small fragments and transferred to serum-free DMEM HG (Sigma-Aldrich) containing collagenase type IV (2 mg/ml, Worthington) and DNase I (20 U/ml, Sigma-Aldrich) (10 ml per lung), and incubated for 30 min at 37°C with agitation (250–260 rpm). Following digestion, samples were centrifuged at 1500 rpm for 5 min at 4°C, and red blood cells were lysed using ACK buffer (Lonza) for 5 min at room temperature. Lysis was quenched with DMEM HG supplemented with 10% FBS, and cells were pelleted by centrifugation. The resulting cell suspension, including residual tissue fragments, was resuspended in complete DMEM HG and plated in 10 cm culture dishes. On day 3, fresh medium was added without disturbing adherent cells. By day 6, cultures reached confluence with proliferating fibroblasts and were used for downstream applications. 2-4 hr prior stimulation medium was replaced to serum-free DMEM HG. The purity of isolated lung fibroblasts we confirmed by flow cytometric analysis with anti-mouse CD140a.

All stimulations and pre-treatments for cells and cell lines are described in the main text and figure legends.

### CRISRP-Cas9 KO of ER-HoxB8 cells

Single guide RNA (sgRNA) lentiviral expression vectors (Vector Builder) contained guides targeting *Rnf128*, *Marchf1*, or non-targeting scramble control and included fluorescent reporters (mCherry or TagBFP2). Lentivirus was produced by transient transfection of HEK293T cells with sgRNA plasmids together with psPAX2 and pMD2.G packaging plasmids using Lipofectamine 3000 (Thermo Fisher Scientific). Viral supernatants were filtered and collected 48 h post-transfection, filtered through 0.45 µm PVDF filters (Milipore Sigma), and used immediately for transduction. Cas9-expressing ER-HoxB8 progenitors were maintained in progenitor medium as described above and cultured at densities below 0.5 × 106 cells/ml. For transduction, progenitors (0.5 × 106 cells per well) were plated in non-tissue culture-treated 6-well plates and incubated with filtered viral supernatant in the presence of polybrene. Spinfection was performed at 1000 × g for 120 min at 26°C, followed by the addition of fresh progenitor medium. Transduced cells were enriched by fluorescence-activated cell sorting based on reporter expression and subsequently expanded for 1–3 weeks while maintaining sub-confluent densities. Cells were either cryopreserved (10% DMSO in FBS) or used for downstream differentiation and genome editing analyses. Target gene disruption was confirmed by Sanger sequencing.

### PA01 culture

Bacteria were cultured by plating the glycerol stock on LB plates. A single colony from the plate was picked and grown overnight in LB broth. Second day, the bacteria were subcultured for 1 hour to bring them to the linear growth phase. The number of bacteria was estimated by measuring the optical density at 600 nm. The numbers were confirmed using standard colony forming unit by plating the inoculum on agar plates.

### Enzyme activity measurements

LasB catalytic activity was measured using SensoLyte Green Elastase Assay Kit (AnaSpec). Briefly, purified LasB +/- phosphoramidon (Cayman Checmial) and Serpins were incubated with natural substrate elastin labeled with the 5-FAM fluorophore and the QXL™ 520 quencher for 30-60 min at RT. Proteolytic cleavage of labeled elastin yielded green fluorescence that was monitored at excitation/emission = 488 nm/520 nm. Increase in fluorescence intensity was directly proportional to LasB activity. Control samples containing only labeled elastin served as blanks.

### *In vitro* cleavage assay

Recombinant mouse Serpine1 and Serpina3n were incubated with LasB for 30-60 min at 37 °C. Reactions were stopped by addition of 4X Laemmli Sample Buffer (200 mM Tris-HCl pH 6.8, 4% SDS, 40% glycerol, 0.4% bromophenol blue, 20 % ß-mercaptoethanol). Reaction products were resolved using SDS-PAGE using 4-15% TGX protein gels (Bio-Rad) and stained by SimplyBlue SafeStain (Themo Fisher Scientific).

### Immunoblotting

Cells were plated and stimulated as indicated in respective figures in 24-well tissue culture plates at densities 2.5 x 10^5^ cells per well. After treatments and stimulations cells were washed twice with ice-cold PBS and lysed in cold buffer consisting of 20 mM Tris (pH 7.5), 150 mM NaCl, 1% SDS, 10 % glycerol, 0.1 % bromophenol blue, 2.5 % ß-mercaptoethanol, 1 × Halt protease and phosphatase inhibitors mix (Thermo Fisher Scientific). Lysates were boiled for 10-15 minutes at 95 °C and stored at -20 °C until further use. Lysates were resolved by SDS -PAGE on 4–15% TGX gradient gels (Bio-rad) and transferred to PVDF Immobilon P membranes (Millipore Sigma) using Trans-Blot Turbo Transfer System (Bio-Rad). Membranes were blocked in TBS-T (tris buffered saline with tween) containing 5% BSA (FisherBio) and probed overnight at 4°C with primary antibodies in TBS-T containing 5% BSA. Fluorophore-conjugated secondary antibodies (anti-mouse Alexa-488 A32723 and anti-rabbit Alexa-800 A32735) were Invitrogen (Thermo Fisher Scientific). Immunoblots were visualized using IamgeLab gel imager (BioRad). Primary antibodies used in this study: phospho-STAT6 (cat#56554), total STAT6 (cat#5397), ß-actin (cat#3700) were from Cell Signaling Technologies.

### RNA isolation and qPCR

Primary BMDMs and lung fibroblasts were plated and stimulated as indicated in respective figures and figure legends in 24-well plates or 12-well plates at densities 2.5 x 10^5^ (for BMDMs) or 0.4 x 10^5^ (for fibroblasts) cells per well. After treatments and stimulations cells were washed twice with ice-cold PBS (Sigma or Thermo Fisher Scientific) and lysed with RNA-Bee or STAT- 60 (Tel-Test) for RNA extraction. RNA extraction was performed according to manufacturer’s instructions. Briefly, 0.2 ml chloroform was added to 0.5-1 ml of RNA-Bee/STAT-60, samples were mixed for 30 seconds, kept on ice for 5 min and centrifuged 12,000g for 15 minutes at 4 °C. 0.5 ml of isopropanol was added to aqueous phase (RNA), samples were kept for 5 min at 4 °C and centrifuged as above. RNA pellet was washed twice with 75% ethanol. The final RNA pellet was air-dried for 10-15 min and resuspended in molecular biology grade water (Sigma). Total RNA was reverse transcribed with an oligo (dT) primer and a Moloney murine leukemia virus reverse transcriptase (MMLV RT, TakaraBio). cDNA was analyzed by quantitat ive PCR amplification using SYBR Green qPCR Master Mix (QuantaBio) or PowerUP SYBR mix (Thermo Fisher Scientific) on a Bio-Rad CFX96 or 384 Real-Time PCR Detection System. Primers were designed to amplify mRNA-specific sequences, and analysis of the melt-curve confirmed the amplification of single products. Unstimulated samples were used as controls. Relative expression was normalized to ribosomal protein L13a (*Rpl13a*). Alternatively, RNA was extracted and purified using RNAeasy mini kit (Qiagen) according to manufacturer’s instructions.

### BAL collection, measurements of total protein, heme and cell counts

Bronchoalveolar lavage fluid (BALF) was collected from euthanized mice by exposing the trachea and cannulating it with a 26-gauge needle fitted with plastic tubing attached to a 1 mL syringe. Lungs were lavaged with 1 mL PBS, which was slowly instilled intratracheally and gently aspirated. Recovered fluid was centrifuged at 2000 rpm for 5 min at 4°C to separate supernatant and cellular fractions. The supernatant was used for biochemical analyses, including total protein quantification using the Pierce BCA Protein Assay Kit (Thermo Fisher Scientific) and heme measurement using a Heme Assay Kit (Sigma-Aldrich), according to the manufacturers’ instructions. Cell pellets were resuspended in 0.3 mL PBS, and total and differential cell counts, including white blood cells, red blood cells, and platelets, as well as hemoglobin and hematocrit levels, were measured using a Coulter Ac·T diff2 Hematology Analyzer (Beckman Coulter). Cells were subsequently pelleted, subjected to red blood cell lysis using ACK lysis buffer (Lonza), washed, and resuspended in 200 µL PBS for downstream flow cytometry analysis.

### Measurements of metabolic analytes

Measurements of metabolites was performed on mouse blood using the Piccolo Xpress Chemistry Analyzer (Abaxis). Whole blood (∼150 µl) was collected via retro-orbital bleeding into lithium heparin-coated tubes. Samples were processed according to the manufacturer’s instructions and 110 µl of blood was loaded onto Piccolo Comprehensive Metabolic Panel (Abbot) reagent discs. Analyses were performed on the Piccolo Xpress system, which automatically quantified standard biochemical parameters, including electrolytes, live r enzymes, renal function markers, and metabolic substrates. Instrument calibration and quality control were performed in accordance with the manufacturer’s guidelines prior to sample analysis.

### Pulmonary function test and analysis

Pulmonary function test was performed using the FlexiVent apparatus (SCIREQ, Montreal, QC, Canada). Specifically, mice were anesthetized with urethane (intraperitoneal injection, 1 -2 g/kg). The mouse was placed supine, and the trachea was cannulated with a 20-gauge tracheostomy tube. The tube was inserted through a small ventral incision made in the rostral-most part of the trachea. The tube was then held securely in place with a suture tied around the trachea. The chest remained closed during all measurements. Mice were mechanically ventilated using the SCIREQ FlexiVent apparatus with 150 breaths/min, a tidal volume of 10 mL/kg body mass, and a positive end-expiratory pressure of 3 cmH2O prior to lung function measurements. Mice were then paralyzed with an intraperitoneal injection of pancuronium bromide (1 mg/kg). With the maximal vital capacity perturbation (called total lung capacity by SCIREQ), the inspiratory capacity (IC) of the lungs was determined using the SCIREQ software (Flexiware v.7.6, Service Pack 6). Forced oscillation perturbations (“quickprime-3”) subsequently measured tissue damping, reflecting energy dissipation within the lung parenchyma. Pressure-volume loops were calculated through quasi-static stepwise pressure-guided measurements of pressure P and volume V. SCIREQ software calculated static compliance (Cst) by fitting the Salazar–Knowles equation V = (A Be^-kP^) to the expiratory portion of the loop and hysteresis. All maneuvers and perturbations were performed until three consistent measurements were achieved per mouse. A coefficient of determination of 0.95 was the lower limit for suitable measurements.

### Micro-computed tomography of lung tissue

Mice were scanned in a supine position using an *in vivo* small animal hybrid SPECT/high-resolution computed tomography (CT) scanner (MI Labs USPECT4/CT). Mice were sedated with isoflurane inhalation via nose cone. Thoracic movement was detected by a motion sensor adhered to the mouse’s thorax allowing division of the respiratory cycle into four phases ranging from maximal inspiration to maximal expiration. Images were obtained throughout the respiratory cycle and sorted to the appropriate phase of respiration during post -acquisition processing. 3D images were constructed using 3D Slicer 4.10.2 r28257 in which aerated regions of lung were segmented from tissue and bone using Houndsfield units (HU) for air as the threshold. The threshold for this study was set to -300 HU. Aerated lung volumes were then calculated.

### Measurements of vital signs

Blood oxygen saturation, breath rate, and heart rate were measured by pulse oximetry using the MouseOx Plus (Starr Life Sciences Corp.). Core body temperature was measured by rectal probe thermometry (Physitemp TH-5 Thermalert).

### Metabolic cage studies

Metabolic phenotyping was performed using a Promethion metabolic cage system (Sable Systems International). Mice were singly housed with ad libitum access to food and water under controlled temperature and a 12 h light–dark cycle. Animals were acclimated to the cages for 24 h followed by three days of steady-state recording before experimentation. Oxygen consumption (VO2), carbon dioxide production (VCO2), respiratory exchange ratio (RER), energy expenditure were continuously measured by indirect calorimetry, while food and water intake and locomotor activity were simultaneously recorded. Data were collected over 96 -72 h and analyzed using the manufacturer’s software.

### Flow cytometry

Mice were euthanized and transcardially perfused with 10 mL PBS supplemented with heparin (10 U mL-¹) using a 26 -gauge needle. The thoracic cavity was rapidly exposed, and the needle was inserted into the right ventricle to selectively perfuse the lungs. Lung were excised, minced into small fragments and transferred to 10 ml of PBS containing collagenase type IV (2 mg/ml, Worthington) and DNase I (20 U/ml, Sigma-Aldrich), and incubated for 30 min at 37°C with agitation (250–260 rpm). Digested tissues were passed through a 70 µm cell strainer into 6-well plates and mechanically dissociated using a syringe plunger, with repeated washes in PBS to maximize cell recovery. Cell suspensions were collected, centrifuged at 2000 rpm for 6 min at 4°C. For BAL samples were processed as described above.

Spun down cell were washed with PBS and stained with Zombie Red Cell Viability Dye (BioLegend) for 10 min at RT. Cell were washed with FACS buffer (PBS supplemented with 2.5 % FBS) and stained with fluorochrome-conjugated antibodies +/- anti-CD16/CD32 (cat#14-0161-86, Thermo Fisher Scientific) diluted in FACS buffer with addition of 123count eBeads (Thermo Fisher Scientific) for 30 min at 4 °C. For intracellular staining, cells were permeabilized and fixed using FoxP3 fixation/permeabilization buffer (Themo Fisher Scientific) following the manufacturer-recommended protocol and incubated with specific fluorochrome-conjugated antibodies for the specified proteins in permeabilization buffer for 30 minutes at 4 °C. Cell acquisition was performed on LSRII instrument (BD Biosciences), and data were analyzed with FlowJo software (Tree Star). Neutrophils were defined as MHCII^−^CD11b^+^ CD64^-^ Ly6G^+^, eosinophils as MHCII^−^ CD11b^+^ CD64^-^ Siglec-F^+^, alveolar macrophages as MHCII^+^ MerTK^+^ CD64^+^ CD11b^-^ SiglecF^+^, interstitial macrophages as MHCII^+^ MerTK^+^ CD64^+^ CD11b^+^ SiglecF^-^, monocyte-derived macrophages as MHCII^+^ MerTK^+^ CD64^+^ CD11b^+^ Ly-6C^+^, monocytes as MHCII^-^ CD64^+^ CD11b^+^ Ly-6C^+^. The following fluorochrome-conjugated antibodies were used: anti-mouse MHCII FITC (I-A/I-E, Thermo Fisher Scientific, cat#11-5321-85), anti-mouse Siglec-F PE (BD Biosciences, cat#552126), anti-mouse Ly-6C APC (Thermo Fisher Scientific, cat#17-5932-82), anti-mouse CD3e V500 (BD Biosciences, cat#560771), anti-mouse CD4 BV 510 (Biolegend, cat#100553), anti-mouse CD8a PerCP/Cy5.5 (BD Biosciences, cat#551162), anti-mouse TCRß APC (Biolegend, cat#109212), anti-mouse TCRγd PE (Thermo Fisher Scientific, cat#MA5-28833), anti-mouse CD3e FITC (Thermo Fisher Scientific, cat#11-0031-82), anti-mouse CD5 FITC (Biolegend, cat#100605), anti-mouse CD8a FITC (Biolegend, cat#100706), anti-mouse CD11b FITC (Thermo Fisher Scientific, cat#11-0112-82), anti-mouse CD11c FITC (Thermo Fisher Scientific, cat#11-0114-82), anti-mouse CD19 FITC (Thermo Fisher Scientific, cat#11-0191-82), anti-mouse CD41 FITC (Biolegend, cat #133903), anti-mouse CD49b FITC (Biolegend, cat#103503), anti-mouse GR-1 FITC (Biolegend, cat#108406), anti-mouse NK1.1 FITC (Biolegend, cat#108706), anti-mouse Ter119 FITC (Biolegend, cat#116206), anti-mouse CD4 FITC (Biolegend, cat#100406), anti-mouse TCRß FITC (Biolegend, cat#109206), anti-mouse CD90.2 BUV737 (BD Biosciences, cat#741702), anti-mouse CD25 BV421 (BD Biosciences, cat#562606), anti-mouse Rorγt PE (Thermo Fisher Scientific, cat#12-6981-82), anti-mouse Gata3 APC (BD Biosciences, cat#567633), anti-mouse CD11b BUV737 (BD Biosciences, cat# 564443), anti-mouse CD45 BUV395 (BD Biosciences, cat# 564279), anti-mouse CD64 Brilliant Violet 421 (BioLegend, cat# 139309), anti-mouse MerTK Pe-Cy7 (Thermo Fisher Scientific, cat#25-5751-82), anti-mouse Ly6G PerCP/Cy5.5 (BD Biosciences, cat # 560602), anti-mouse CD326 Pe-Cy7 (Biolegend, cat #118215), anti-mouse CD140a APC (Thermo Fisher Scientific, cat#17 -1401-81), anti-mouse CD31 PE (Thermo Fisher Scientific, cat#14-0311-82).

### Cytokine ELISA

Cytokine levels in BALF and plasma were quantified using a multiplex bead-based immunoassay (Luminex). BAL supernatants and plasma samples were collected, clarified by centrifugation to remove cellular debris, and stored at -20° C. Samples were processed according to the manufacturer’s instructions using commercially available multiplex panels. Briefly, analytes were captured using fluorescently coded microspheres conjugated to target -specific antibodies, followed by detection with biotinylated secondary antibodies and streptavidin - phycoerythrin. Data were acquired on a Luminex platform, and cytokine conc entrations were determined from standard curves generated using recombinant protein standards. Instrument calibration and quality control were performed according to the manufacturer’s guidelines.

### Lung histology

Lungs were perfused through the right ventricle with 10 ml PBS, removed, and immersed in 4% PFA or 10% formalin for 24-96 h, followed by 70% ethanol until embedding in paraffin. Tissues were sliced, and 5-mm sections were stained with H&E, Verhoeff Van Giesen (elastin stain), Masson’s trichrome (collagen stain) or Movat pentachrome (stain for collagen, glycosaminoglycans, muscle, elastic fibers and nuclei). 5 -10 images per mouse per experimental conditions were used for quantification.

#### Quantification of cell numbers

Images were opened in Fiji (ImageJ) and processed using “Color Deconvolution” plugin with selection of specific type of staining: H&E. Images were deconvoluted in their green, red, and blue components. The red component was further used for measurements of cell numbers. The area of the red image was measured using the “Threshold” tool. The threshold value was manually adjusted to modify the selected area. The same value of the threshold was applied to all processed images. Binary image generated after the application of the threshold value was processed using the “Watershed” function to separate individual cells. Next, the image was analyzed using “Analyze Particles” tool. The minimum particle size was set to 0.01 mm^2^ and the “Outline” was selected in the show section. The counted cells with their total area were generated as “Summary” and “Results” measurements. The total image area was quantified to measure cell density per area.

#### Quantification of collagen and elastin

Following background subtraction and pixel-based thresholding, area fractions for collagen (from Masson’s Trichrome), elastin (from VVG) and cytoplasm (from MTC) were computed as the ratio of pixels corresponding to a stain divided by the total number of pixels in the image. Collagen area fraction was computed as 1 - area fraction of elastin plus cytoplasm. Glycosaminoglycan content was assumed to be negligible as confirmed with Movat Pentachrome staining.

#### Immunofluorescence and quantification

Lung tissues were perfused and fixed in 1% paraformaldehyde overnight at 4°C. Tissues were washed with 5mM NH4Cl and incubated in 30% (w/v) sucrose overnight at 4°C. Tissues were embedded in Optimal Cutting Temperature Compound (Tissue-Tek), flash-frozen, and stored at -80° C. 10 µm sections were cut on CM1950 Cryostat (Leica, Wetzlar, Germany) and fixed in ice-cold acetone for 7 minutes, blocked with 1% rat serum/1% mouse serum and 1% Fc Block and stained with CD169-BUV594 (5µg/mL) (Biolegend, cat#142416), Vimentin-AF647 (5µg/mL) (Biolegend, cat#677807), Hoechst 33,342 (2µg/mL) (Invitrogen Waltham, MA cat#H3570) and sealed with ProLong Gold Antifade (Invitrogen, cat#P36930). Images were captured using a Leica Stellaris 8 Confocal Microscope. Fluorescent channels were photographed separately and then merged. Exposure times and fluorescence intensities were normalized to appropriate control isotype images.

#### Quantification of cell numbers and distance

Macrophage and fibroblast numbers were quantified from confocal images using Fiji. Regions of interest (ROIs) were manually defined and applied to restrict analysis to the relevant whole lung lobe. Images were preprocessed by conversion to 8-bit, followed by median filtering (radius = 2) and Gaussian blurring (s = 2) to reduce noise. Automated thresholding was performed using the Otsu method, and images were converted to binary masks. Segmentation of adjacent cells was improved using the watershed algorithm. Macrophages were identified and quantified using the “Analyze Particles” function with size thresholds of 80-500 µM² and circularity parameters of 0.5-1.0. Fibroblasts were quantified with size thresholds of 75-500 µM² and circularity parameters of 0.2-0.8. Total cell counts and ROI areas were recorded to measure cell density per area. Image calibration was applied with 1792 pixels corresponding to 1000 µm to calculate ROI area in mm². Cell densities were calculated as the number of cells per mm². To assess spatial relationships between macrophages and fibroblasts, fibroblast and macrophage images were processed sequentially as above. Centroid coordinates for each identified cell were extracted using the “Analyze Particles” function. Pairwise Euclidean distances between macrophage and fibroblast centroids were subsequently calculated in Microsoft Excel, and the minimal distance between each macrophage and its nearest fibroblast neighbor was determined.

### Bulk lung RNA-seq and analysis

RNA was extracted from samples using RNeasy Mini Kit with on-column DNase digestion, following manufacturer’s instructions (Qiagen, 74904). RNA quality and integrity were assessed using the Agilent 4200 TapeStation, with RNA integrity numbers (RIN) = 8 considered acceptable. RNA-seq libraries were prepared using Illumina TruSeq Stranded mRNA kit following the manufacturer’s protocol. Library concentration was measured using the KAPA RNA HyperPrep according to manufacturer’s instructions (Roche, 08098131702). RNA-seq libraries were pooled and sequenced on an Illumina NovaSeq 6000 (2 × 100 bp). Reads were mapped to mouse genome GRCm39 and counts were normalized to transcripts per million (TPM) using “kallisto” version 0.48.0 (*101*). Pairwise differential gene expression using Likelihood Ratio Test (LRT) was performed with “DESeq2” version 1.42.0 (*102*). Transcription factor regulatory network and metabolic pathway enrichment analyses were performed on significantly differentially expressed genes (adjusted P < 0.05, Log2FC > 1) using the TRRUST and MouseCyc databases, respectively, through the Metascape and ShinyGO platforms. Enriched terms were considered statistically significant at a false discovery rate (FDR)-adjusted P value < 0.05.

### Lung scRNA-seq and analysis

#### Library preparation and sequencing

Single-cell suspensions were prepared from freshly isolated lung tissue and filtered through a 70-µm cell strainer to remove debris and aggregates. Cell viability was assessed by trypan blue exclusion and samples with viability >80% were used for library preparation. Cells were resuspended in PBS containing 0.04% BSA at the concentration of 1000 cells/µl. Single-cell libraries from 10,000 cells per sample were prepared using the Chromium Single Cell 3’ Gene Expression kit v3.1 (10x Genomics) following the manufacturer’s protocol. Final libraries were quantified using Qubit fluorometric quantification (ThermoFisher Scientific) and quality was assessed using the Agilent 4200 TapeStation. The generated scRNA-seq libraries were sequenced on a NovaSeq 6000 (Illumina) with paired-end (2 × 100 bp). Sequencing was performed at a depth of at least 20,000 read pairs per cell.

#### Data processing and analysis

Raw sequencing reads were aligned to the mouse reference genome (mm10) and gene-cell count matrices were generated using CellRanger (7.1.0). Unfiltered CellRanger outputs were imported into R as Seurat objects (v5.1.0) (*103*), and empty droplets were excluded using a UMI rank cutoff of 30,000 for all samples, with the exception of one superloaded sample (LasB1d_3), for which a rank cutoff of 40,000 was applied. Droplets were retained if they contained a minimum of 125 UMI counts and 125 detected features, and less than 15% mitochondrial reads. Following initial filtration, additional low-quality cells and potential doublets were removed through cluster-based cleaning, in which clusters exhibiting uniquely low information or co-expression of distinct lineage markers were excluded. Epithelial cells were removed for subsequent analyses due to low cell viability. Clean sample objects were merged into a single Seurat object and normalized using log normalization. Dimensionality reduction was performed by principal component analysis (50 PCs) followed by UMAP embedding and unsupervised Louvain clustering (0.5 resolution). Cell classes and cell types were annotated based on expression of canonical marker genes. Cell-cell communication analysis was performed using CellChat (v2.2) (*104*) to infer intercellular signaling interactions across merged immune, mesenchymal, and endothelial compartments. Custom plots were generated using the following R packages: ggplot2 (v3.5.1), viridis (v0.6.5), ComplexHeatmap (v2.22.0) (*105*), and cowplot (v1.2.0). For data processing and analysis code, see our Github repository. Cell -cell communication was inferred using CellChat (v2.2) (*104*), with communication probabilities computed via the trimean method (computeCommunProb, type = “triMean”) and restricted to cell groups containing =10 cells. Differential gene expression analysis was performed on scRNA-seq data separately within immune and mesenchymal cell populations of the PBS condition using Seurat’s FindAllMarkers function with the Wilcoxon rank-sum test. Transcription factor regulatory network and metabolic pathway enrichment analyses were performed as described above for bulk lung RNA-seq analysis. Analyses of gene expression in public scRNA-seq data sets (*62*, *63*) were performed using the interactive Single Cell Portal – Broad Institute (https://singlecell.broadinstitute.org/single_cell) and COPD Cell Atlas (https://www.copdcellatlas.com/).

### Quantification and statistical analysis

Sample or experiment sizes were determined empirically for statistical power. No statistical tests were used to predetermine the size of experiments. Statistical tests were performed using GraphPad Prism, with significance defined as p =0.05. Statistical identification of outliers was performed using either the ROUT (Robust regression and Outlier removal) method or Grubbs’ test. ROUT analysis was conducted with a false discovery rate (Q) of 1%, which allows robust detection of extreme values while controlling for type I error. Grubbs’ test was applied to datasets with a single suspected outlier and normal distribution, using an alpha level of 0.05. For two group comparisons, an unpaired Student’s t test was used. More than two groups were compared using one-way analysis of variance (ANOVA). To perform post hoc comparison of selected group means we performed Bonferroni’s test. Two-way ANOVA was conducted to examine the combined and individual impact of two independent variables. The log-rank Mantel-Cox test was used to compare survival curves between groups.

## References and Notes

1. R. Chovatiya, R. Medzhitov, Stress, inflammation, and defense of homeostasis. Mol Cell 54, 281–288 (2014).

2. D. Kultz, Molecular and evolutionary basis of the cellular stress response. Annu Rev Physiol 67, 225–257 (2005).

3. M. L. Meizlish, R. A. Franklin, X. Zhou, R. Medzhitov, Tissue Homeostasis and Inflammation. Annu Rev Immunol 39, 557–581 (2021).

4. M. Costa-Mattioli, P. Walter, The integrated stress response: From mechanism to disease. Science 368, (2020).

5. S. Solanki, Y. M. Shah, Hypoxia-Induced Signaling in Gut and Liver Pathobiology. Annu Rev Pathol 19, 291–317 (2024).

6. E. B. Kopp, K. Agaronyan, I. Licona-Limon, S. A. Nish, R. Medzhitov, Modes of type 2 immune response initiation. Immunity 56, 687–694 (2023).

7. M. E. Kotas, R. Medzhitov, Homeostasis, inflammation, and disease susceptibility. Cell 160, 816–827 (2015).

8. R. Medzhitov, The spectrum of inflammatory responses. Science 374, 1070–1075 (2021).

9. C. Lopez-Otin, J. S. Bond, Proteases: multifunctional enzymes in life and disease. J Biol Chem 283, 30433–30437 (2008).

10. X. S. Puente, L. M. Sanchez, C. M. Overall, C. Lopez-Otin, Human and mouse proteases: a comparative genomic approach. Nat Rev Genet 4, 544–558 (2003).

11. S. S. Twining, Regulation of proteolytic activity in tissues. Crit Rev Biochem Mol Biol 29, 315–383 (1994).

12. A. P. Amini et al., Multiscale profiling of protease activity in cancer. Nat Commun 13, 5745 (2022).

13. P. J. Barnes et al., Chronic obstructive pulmonary disease. Nat Rev Dis Primers 1, 15076 (2015).

14. S. S. Glasson et al., Deletion of active ADAMTS5 prevents cartilage degradation in a murine model of osteoarthritis. Nature 434, 644–648 (2005).

15. S. M. McCarty, S. L. Percival, Proteases and Delayed Wound Healing. Adv Wound Care (New Rochelle) 2, 438–447 (2013).

16. A. Janoff, J. D. Zeligs, Vascular injury and lysis of basement membrane in vitro by neutral protease of human leukocytes. Science 161, 702–704 (1968).

17. L. Thomas, Reversible collapse of rabbit ears after intravenous papain, and prevention of recovery by cortisone. J Exp Med 104, 245–252 (1956).

18. B. W. Zweifach, A. L. Nagler, W. Troll, Some effects of proteolytic inhibitors on tissue injury and systemic anaphylaxis. J Exp Med 113, 437–450 (1961).

19. S. J. Galli, M. Tsai, Mast cells: versatile regulators of inflammation, tissue remodeling, host defense and homeostasis. J Dermatol Sci 49, 7–19 (2008).

20. P. Konieczny et al., Interleukin-17 governs hypoxic adaptation of injured epithelium. Science 377, eabg9302 (2022).

21. S. Naik et al., Inflammatory memory sensitizes skin epithelial stem cells to tissue damage. Nature 550, 475–480 (2017).

22. J. Zhao, I. Andreev, H. M. Silva, Resident tissue macrophages: Key coordinators of tissue homeostasis beyond immunity. Sci Immunol 9, eadd1967 (2024).

23. K. Agaronyan et al., Tissue remodeling by an opportunistic pathogen triggers allergic inflammation. Immunity 55, 895–911 e810 (2022).

24. E. Florsheim et al., Integrated innate mechanisms involved in airway allergic inflammation to the serine protease subtilisin. J Immunol 194, 4621–4630 (2015).

25. R. K. Rosenstein, J. S. Bezbradica, S. Yu, R. Medzhitov, Signaling pathways activated by a protease allergen in basophils. Proc Natl Acad Sci U S A 111, E4963–4971 (2014).

26. C. L. Sokol, G. M. Barton, A. G. Farr, R. Medzhitov, A mechanism for the initiation of allergen-induced T helper type 2 responses. Nat Immunol 9, 310–318 (2008).

27. A. Babtie, N. Tokuriki, F. Hollfelder, What makes an enzyme promiscuous? Curr Opin Chem Biol 14, 200–207 (2010).

28. J. G. Walsh, S. E. Logue, A. U. Luthi, S. J. Martin, Caspase-1 promiscuity is counterbalanced by rapid inactivation of processed enzyme. J Biol Chem 286, 32513–32524 (2011).

29. C. M. Greene et al., alpha1-Antitrypsin deficiency. Nat Rev Dis Primers 2, 16051 (2016).

30. M. Meyer, I. Jaspers, Respiratory protease/antiprotease balance determines susceptibility to viral infection and can be modified by nutritional antioxidants. Am J Physiol Lung Cell Mol Physiol 308, L1189–1201 (2015).

31. C. Taggart et al., Protean proteases: at the cutting edge of lung diseases. Eur Respir J 49, (2017).

32. H. Aegerter, B. N. Lambrecht, C. V. Jakubzick, Biology of lung macrophages in health and disease. Immunity 55, 1564–1580 (2022).

33. E. I. Arafa, et al., Recruitment and training of alveolar macrophages after pneumococcal pneumonia. JCI Insight 7, (2022).

34. H. E. Ghoneim, P. G. Thomas, J. A. McCullers, Depletion of alveolar macrophages during influenza infection facilitates bacterial superinfections. J Immunol 191, 1250–1259 (2013).

35. F. Li, et al., Monocyte-derived alveolar macrophages autonomously determine severe outcome of respiratory viral infection. Sci Immunol 7, eabj5761 (2022).

36. A. V. Misharin et al., Monocyte-derived alveolar macrophages drive lung fibrosis and persist in the lung over the life span. J Exp Med 214, 2387–2404 (2017).

37. S. Verwaerde et al., Innate type 2 lymphocytes trigger an inflammatory switch in alveolar macrophages. Immunity 59, 60–78 e69 (2026).

38. H. Aegerter et al., Influenza-induced monocyte-derived alveolar macrophages confer prolonged antibacterial protection. Nat Immunol 21, 145–157 (2020).

39. Y. Wang et al., Eosinophils attenuate hepatic ischemia-reperfusion injury in mice through ST2-dependent IL-13 production. Sci Transl Med 13, (2021).

40. B. G. Yipp, et al., The Lung is a Host Defense Niche for Immediate Neutrophil-Mediated Vascular Protection. Sci Immunol 2, (2017).

41. C. Yu et al., Targeted deletion of a high-affinity GATA-binding site in the GATA-1 promoter leads to selective loss of the eosinophil lineage in vivo. J Exp Med 195, 1387–1395 (2002).

42. M. Koupenova, L. Clancy, H. A. Corkrey, J. E. Freedman, Circulating Platelets as Mediators of Immunity, Inflammation, and Thrombosis. Circ Res 122, 337–351 (2018).

43. A. T. Nurden, Platelets, inflammation and tissue regeneration. Thromb Haemost 105 Suppl 1, S13–33 (2011).

44. N. V. Serbina, E. G. Pamer, Monocyte emigration from bone marrow during bacterial infection requires signals mediated by chemokine receptor CCR2. Nat Immunol 7, 311–317 (2006).

45. P. Mombaerts et al., RAG-1-deficient mice have no mature B and T lymphocytes. Cell 68, 869–877 (1992).

46. E. Camerer, W. Huang, S. R. Coughlin, Tissue factor- and factor X-dependent activation of protease-activated receptor 2 by factor VIIa. Proc Natl Acad Sci U S A 97, 5255–5260 (2000).

47. M. Steinhoff et al., Agonists of proteinase-activated receptor 2 induce inflammation by a neurogenic mechanism. Nat Med 6, 151–158 (2000).

48. L. F. Loffredo et al., An amphiregulin reporter mouse enables transcriptional and clonal expansion analysis of reparative lung Tregs. JCI Insight 10, (2025).

49. C. M. Minutti et al., A Macrophage-Pericyte Axis Directs Tissue Restoration via Amphiregulin-Induced Transforming Growth Factor Beta Activation. Immunity 50, 645–654 e646 (2019).

50. P. Baral et al., Nociceptor sensory neurons suppress neutrophil and gammadelta T cell responses in bacterial lung infections and lethal pneumonia. Nat Med 24, 417–426 (2018).

51. Y. Z. Lu et al., CGRP sensory neurons promote tissue healing via neutrophils and macrophages. Nature 628, 604–611 (2024).

52. J. Uddin et al., CGRP-related neuropeptide adrenomedullin 2 promotes tissue-protective ILC2 responses and limits intestinal inflammation. Nat Immunol 26, 1516–1526 (2025).

53. H. Han et al., TRRUST v2: an expanded reference database of human and mouse transcriptional regulatory interactions. Nucleic Acids Res 46, D380–D386 (2018).

54. K. Chan, Y. W. Kan, Nrf2 is essential for protection against acute pulmonary injury in mice. Proc Natl Acad Sci U S A 96, 12731–12736 (1999).

55. Y. Ishii et al., Transcription factor Nrf2 plays a pivotal role in protection against elastase-induced pulmonary inflammation and emphysema. J Immunol 175, 6968–6975 (2005).

56. Q. Ma, Role of nrf2 in oxidative stress and toxicity. Annu Rev Pharmacol Toxicol 53, 401–426 (2013).

57. A. C. Rothchild, et al., Alveolar macrophages generate a noncanonical NRF2-driven transcriptional response to Mycobacterium tuberculosis in vivo. Sci Immunol 4, (2019).

58. M. P. Soares, A. M. Ribeiro, Nrf2 as a master regulator of tissue damage control and disease tolerance to infection. Biochem Soc Trans 43, 663–668 (2015).

59. V. L. Ngo et al., Segmented filamentous bacteria reprogramming of alveolar macrophages limits postinfluenza bacterial pneumonia. Sci Immunol 11, eadt8858 (2026).

60. H. Zhang et al., AMFR drives allergic asthma development by promoting alveolar macrophage-derived GM-CSF production. J Exp Med 219, (2022).

61. M. Guilliams et al., Alveolar macrophages develop from fetal monocytes that differentiate into long-lived cells in the first week of life via GM-CSF. J Exp Med 210, 1977–1992 (2013).

62. M. L. Caton, M. R. Smith-Raska, B. Reizis, Notch-RBP-J signaling controls the homeostasis of CD8- dendritic cells in the spleen. J Exp Med 204, 1653–1664 (2007).

63. B. E. Clausen, C. Burkhardt, W. Reith, R. Renkawitz, I. Forster, Conditional gene targeting in macrophages and granulocytes using LysMcre mice. Transgenic Res 8, 265–277 (1999).

64. L. van de Laar et al., Yolk Sac Macrophages, Fetal Liver, and Adult Monocytes Can Colonize an Empty Niche and Develop into Functional Tissue-Resident Macrophages. Immunity 44, 755–768 (2016).

65. W. X. Zhang et al., A Functional Assessment of Fetal Liver and Monocyte-Derived Macrophages in the Lung Alveolar Environment. J Immunol 212, 1012–1021 (2024).

66. T. Fabre, et al., Identification of a broadly fibrogenic macrophage subset induced by type 3 inflammation. Sci Immunol 8, eadd8945 (2023).

67. A. Jaiswal et al., Spatial transcriptomics reveals altered communities and drivers of aberrant epithelia and pro-fibrotic fibroblasts in interstitial lung diseases. Cell Genom, 101066 (2026).

68. M. Sauler et al., Characterization of the COPD alveolar niche using single-cell RNA sequencing. Nat Commun 13, 494 (2022).

69. F. Chen et al., An essential role for TH2-type responses in limiting acute tissue damage during experimental helminth infection. Nat Med 18, 260–266 (2012).

70. R. L. Gieseck, 3rd, M. S. Wilson, T. A. Wynn, Type 2 immunity in tissue repair and fibrosis. Nature reviews. Immunology 18, 62–76 (2018).

71. J. A. Knipper et al., Interleukin-4 Receptor alpha Signaling in Myeloid Cells Controls Collagen Fibril Assembly in Skin Repair. Immunity 43, 803–816 (2015).

72. E. H. Kobayashi et al., Nrf2 suppresses macrophage inflammatory response by blocking proinflammatory cytokine transcription. Nat Commun 7, 11624 (2016).

73. L. G. Bankova, et al., The cysteinyl leukotriene 3 receptor regulates expansion of IL-25-producing airway brush cells leading to type 2 inflammation. Sci Immunol 3, (2018).

74. C. Schneider, C. E. O’Leary, R. M. Locksley, Regulation of immune responses by tuft cells. Nature reviews. Immunology 19, 584–593 (2019).

75. M. S. Nadjsombati et al., Detection of Succinate by Intestinal Tuft Cells Triggers a Type 2 Innate Immune Circuit. Immunity 49, 33–41 e37 (2018).

76. J. Oh et al., MARCH1-mediated MHCII ubiquitination promotes dendritic cell selection of natural regulatory T cells. J Exp Med 210, 1069–1077 (2013).

77. A. Sahoo, A. Alekseev, L. Obertas, R. Nurieva, Grail controls Th2 cell development by targeting STAT6 for degradation. Nat Commun 5, 4732 (2014).

78. A. W. Roberts et al., Cas9(+) conditionally-immortalized macrophages as a tool for bacterial pathogenesis and beyond. Elife 8, (2019).

79. M. C. Bouton et al., The under-appreciated world of the serpin family of serine proteinase inhibitors. EMBO Mol Med 15, e17144 (2023).

80. A. Sanchez-Navarro, I. Gonzalez-Soria, R. Caldino-Bohn, N. A. Bobadilla, An integrative view of serpins in health and disease: the contribution of SerpinA3. Am J Physiol Cell Physiol 320, C106–C118 (2021).

81. M. P. Soares, I. Hamza, Macrophages and Iron Metabolism. Immunity 44, 492–504 (2016).

82. P. G. Gettins, Serpin structure, mechanism, and function. Chem Rev 102, 4751–4804 (2002).

83. M. Divangahi et al., Trained immunity, tolerance, priming and differentiation: distinct immunological processes. Nat Immunol 22, 2–6 (2021).

84. F. A. DeLano, D. B. Hoyt, G. W. Schmid-Schonbein, Pancreatic digestive enzyme blockade in the intestine increases survival after experimental shock. Sci Transl Med 5, 169ra111 (2013).

85. L. Deng et al., S. aureus drives itch and scratch-induced skin damage through a V8 protease-PAR1 axis. Cell 186, 5375–5393 e5325 (2023).

86. C. J. Harvey et al., Targeting Nrf2 signaling improves bacterial clearance by alveolar macrophages in patients with COPD and in a mouse model. Sci Transl Med 3, 78ra32 (2011).

87. L. Hecker et al., Reversal of persistent fibrosis in aging by targeting Nox4-Nrf2 redox imbalance. Sci Transl Med 6, 231ra247 (2014).

88. J. M. Marzec et al., Functional polymorphisms in the transcription factor NRF2 in humans increase the risk of acute lung injury. FASEB J 21, 2237–2246 (2007).

89. M. Guilliams, G. R. Thierry, J. Bonnardel, M. Bajenoff, Establishment and Maintenance of the Macrophage Niche. Immunity 52, 434–451 (2020).

90. D. Hashimoto et al., Tissue-resident macrophages self-maintain locally throughout adult life with minimal contribution from circulating monocytes. Immunity 38, 792–804 (2013).

91. M. Haldar et al., Heme-mediated SPI-C induction promotes monocyte differentiation into iron-recycling macrophages. Cell 156, 1223–1234 (2014).

92. G. Kovtunovych, M. A. Eckhaus, M. C. Ghosh, H. Ollivierre-Wilson, T. A. Rouault, Dysfunction of the heme recycling system in heme oxygenase 1 -deficient mice: effects on macrophage viability and tissue iron distribution. Blood 116, 6054–6062 (2010).

93. M. Pfefferle et al., Hemolysis transforms liver macrophages into antiinflammatory erythrophagocytes. J Clin Invest 130, 5576–5590 (2020).

94. A. Hedblom et al., Heme detoxification by heme oxygenase-1 reinstates proliferative and immune balances upon genotoxic tissue injury. Cell Death Dis 10, 72 (2019).

95. F. Vallelian et al., Proteasome inhibition and oxidative reactions disrupt cellular homeostasis during heme stress. Cell Death Differ 22, 597–611 (2015).

96. T. Tsukui, P. J. Wolters, D. Sheppard, Alveolar fibroblast lineage orchestrates lung inflammation and fibrosis. Nature 631, 627–634 (2024).

97. J. Meixiong et al., Identification of a bilirubin receptor that may mediate a component of cholestatic itch. Elife 8, (2019).

98. D. E. Stec et al., Bilirubin Binding to PPARalpha Inhibits Lipid Accumulation. PLoS One 11, e0153427 (2016).

99. M. Lindman et al., Astrocytic RIPK3 exerts protective anti-inflammatory activity in mice with viral encephalitis by transcriptional induction of serpins. Sci Signal 18, eadq6422 (2025).

100. L. Vicuna et al., The serine protease inhibitor SerpinA3N attenuates neuropathic pain by inhibiting T cell-derived leukocyte elastase. Nat Med 21, 518–523 (2015).

101. N. L. Bray, H. Pimentel, P. Melsted, L. Pachter, Near-optimal probabilistic RNA-seq quantification. Nat Biotechnol 34, 525–527 (2016).

102. M. I. Love, W. Huber, S. Anders, Moderated estimation of fold change and dispersion for RNA-seq data with DESeq2. Genome Biol 15, 550 (2014).

103. Y. Hao et al., Dictionary learning for integrative, multimodal and scalable single-cell analysis. Nat Biotechnol 42, 293–304 (2024).

104. S. Jin, M. V. Plikus, Q. Nie, CellChat for systematic analysis of cell-cell communication from single-cell transcriptomics. Nat Protoc 20, 180–219 (2025).

105. Z. Gu, Complex heatmap visualization. Imeta 1, e43 (2022).

